# Minimal requirements for a neuron to co-regulate many properties and the implications for ion channel correlations and robustness

**DOI:** 10.1101/2020.12.04.410787

**Authors:** Jane Yang, Husain Shakil, Stéphanie Ratté, Steven A. Prescott

## Abstract

Neurons regulate their excitability by adjusting their ion channel levels. Degeneracy – achieving equivalent outcomes (excitability) using different solutions (channel combinations) – facilitates this regulation by enabling a disruptive change in one channel to be offset by compensatory changes in other channels. But neurons must co-regulate many properties. Pleiotropy – the impact of one channel on more than one property – complicates regulation because a compensatory ion channel change that restores one property to its target value often disrupts other properties. How then does a neuron simultaneously regulate multiple properties? Here we demonstrate that of the many channel combinations producing the target value for one property (the single-output solution set), few combinations produce the target value for other properties. Combinations producing the target value for two or more properties (the multi-output solution set) correspond to the intersection between single-output solution sets. Properties can be effectively co-regulated only if the number of adjustable channels (*n*_in_) exceeds the number of regulated properties (*n*_out_). Ion channel correlations emerge during homeostatic regulation when the dimensionality of solution space (*n*_in_ – *n*_out_) is low. Even if each property can be regulated to its target value when considered in isolation, regulation as a whole fails if single-output solution sets do not intersect. Our results also highlight that ion channels must be co-adjusted with different ratios to regulate different properties, which suggests that each error signal drives modulatory changes independently, despite those changes ultimately affecting the same ion channels.

## INTRODUCTION

Neurons maintain their average firing rate near a target value, or set point, by adjusting their intrinsic excitability and synaptic weights^1–6^. Homeostatic regulation of intrinsic excitability is achieved through feedback control of diverse ion channels^1, 2, 4, 7^. Computational models have successfully employed negative feedback to adjust ion channel densities^8–12^ and control theory provides a valuable framework to conceptualize how this occurs^13^. But most of the mechanistic details remain unclear^14, 15^ and are not straightforward; for instance, different perturbations can trigger similar changes in excitability via different signaling pathways affecting different ion channels^16^, or via different combinations of excitability changes and synaptic scaling^17^.

Firing rate homeostasis is facilitated by the ability of different ion channel combinations to produce equivalent excitability^18^. The ability of distinct elements to produce the same outcome is known as degeneracy^19^ and has attracted increasing attention in neuroscience^18, 20–27^. Many aspects of neural function at the genetic^28, 29^, synaptic^30, 31^, cellular^32–39^, and network^40–44^ levels are now recognized as being degenerate. Ion channel degeneracy facilitates robust homeostatic regulation of neuronal excitability by enabling a disruptive change in one ion channel to be offset by compensatory changes in other ion channels^22, 34, 35, 45–50^.

But to maintain good neural coding, neurons must regulate various aspects of their activity beyond average firing rate^51^, and must do so while also regulating their energy usage, osmolarity, pH, protein levels, etc.^52–54^. These other cellular properties affect and are affected by neuronal activity, which is to say that properties are not regulated in isolation from one another. And just as each property depends on multiple ion channels, each ion channel can affect multiple properties^33, 38^. This pleiotropy or functional overlap^55^ confounds the independent regulation of each property. In response to a perturbation, a compensatory ion channel change that restores one property to its target value may exacerbate rather than mitigate the disturbance of a second property (**Fig. 1A**). In theory, both properties could be regulated by adjusting two or more ion channels (**Fig. 1B**), but this suggests that the number of properties that can be co-regulated is limited by the number of adjustable ion channels, even if the channels are pleiotropic.

**Figure 1.**
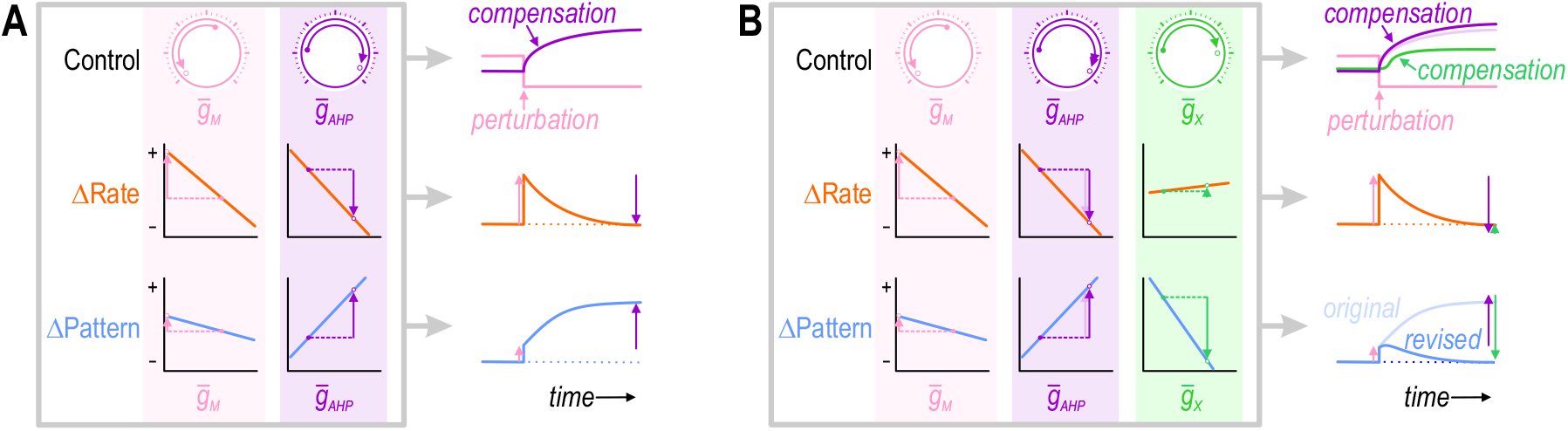
Simultaneous regulation of >1 property in a theoretical neuron. **(A)** Challenge. If two ion channels, *g*_M_ (pink) and *g*_AHP_ (purple), both affect firing rate (orange), then the change in firing rate caused by perturbing *g*_M_ can be offset by a compensatory change in *g*_AHP_. However, if *g*_M_ and *g*_AHP_ also affect firing pattern (blue), but not in exactly the same way, then the compensatory change in *g*_AHP_ that restores firing rate to its target value may exacerbate rather than resolve the change in firing pattern. **(B)** Solution. Adjusting *g*_AHP_ and at least one additional conductance, *g*_x_ (green), that affects each property in a different way than *g*_AHP_, may enable firing rate and firing pattern to be regulated back to their target values. This suggests that the number of adjustable ion channels (“dials”) relative to the number of regulated properties is important. Note that adjusting *g*_x_ to restore the firing pattern necessitates a small extra increase in *g*_AHP_ (compare pale and dark purple); in other words, conductances must be co-adjusted.

To identify the requirements for co-regulating many properties, we began with an experiment to confirm that regulating one property by adjusting a single channel is liable to disrupt other properties. We then proceeded with modeling to unravel how multiple channels must be co-adjusted to ensure proper regulation of >1 property. We show that the number of adjustable ion channels (*n*_in_) must exceed the number of regulated properties (*n*_out_). This is consistent with past work^11, 50, 56^ but the implications for homeostatic regulation have not been explored. To that end, we show that the dimensionality of solution space (*n*_in_ – *n*_out_) influences the emergence of ion channel correlations in the presence or absence of noise, and that increased correlations presage regulation failure. We also show that regulating different properties requires that ion channels are co-adjusted with different ratios, which necessitates separate master regulators.

## RESULTS

### Adjusting an ion channel to regulate one property risks disrupting other properties – experiments

To test experimentally if regulating one property by adjusting one ion channel is liable to disrupt a second property, we disrupted the firing rate of a CA1 pyramidal neuron by blocking its voltage-gated M-type K^+^ current (*I*_M_) and then we restored firing rate to its baseline value by inserting a virtual calcium-activated AHP-type K^+^ current (*I*_AHP_) using dynamic clamp, while also monitoring the firing pattern. Specifically, the neuron was stimulated by injecting irregularly fluctuating (noisy) current under four conditions (**Fig. 2A,B**): at baseline (blue), after blocking native *I*_M_ (black), and again after introducing virtual *I*_M_ (cyan) or virtual *I*_AHP_ (red). Adding virtual *I*_M_ demonstrates that we can replace a native current with an equivalent virtual current; compensation was modeled by inserting a distinct current with functional overlap, namely *I*_AHP_ (see Fig. 1). Inserting virtual *I*_M_ or virtual *I*_AHP_ reversed the depolarization (**Fig. 2C**), spike amplitude attenuation (**Fig. 2D**) and firing rate increase (**Fig. 2E**) caused by blocking native *I*_M_. Replacing native *I*_M_ with virtual *I*_M_ did not affect firing pattern, quantified as the coefficient of variation of the interspike interval (CV_ISI_), whereas virtual *I*_AHP_ reduced CV_ISI_ (**Fig. 2F**), often causing the neuron to spike at different times (in response to different stimulus fluctuations) than the neuron with native or virtual *I*_M_ (**Fig. 2G**). The results confirm predictions from Figure 1, namely that compensatory “upregulation” of *I*_AHP_ restored firing rate but disrupted firing pattern.

**Figure 2.**
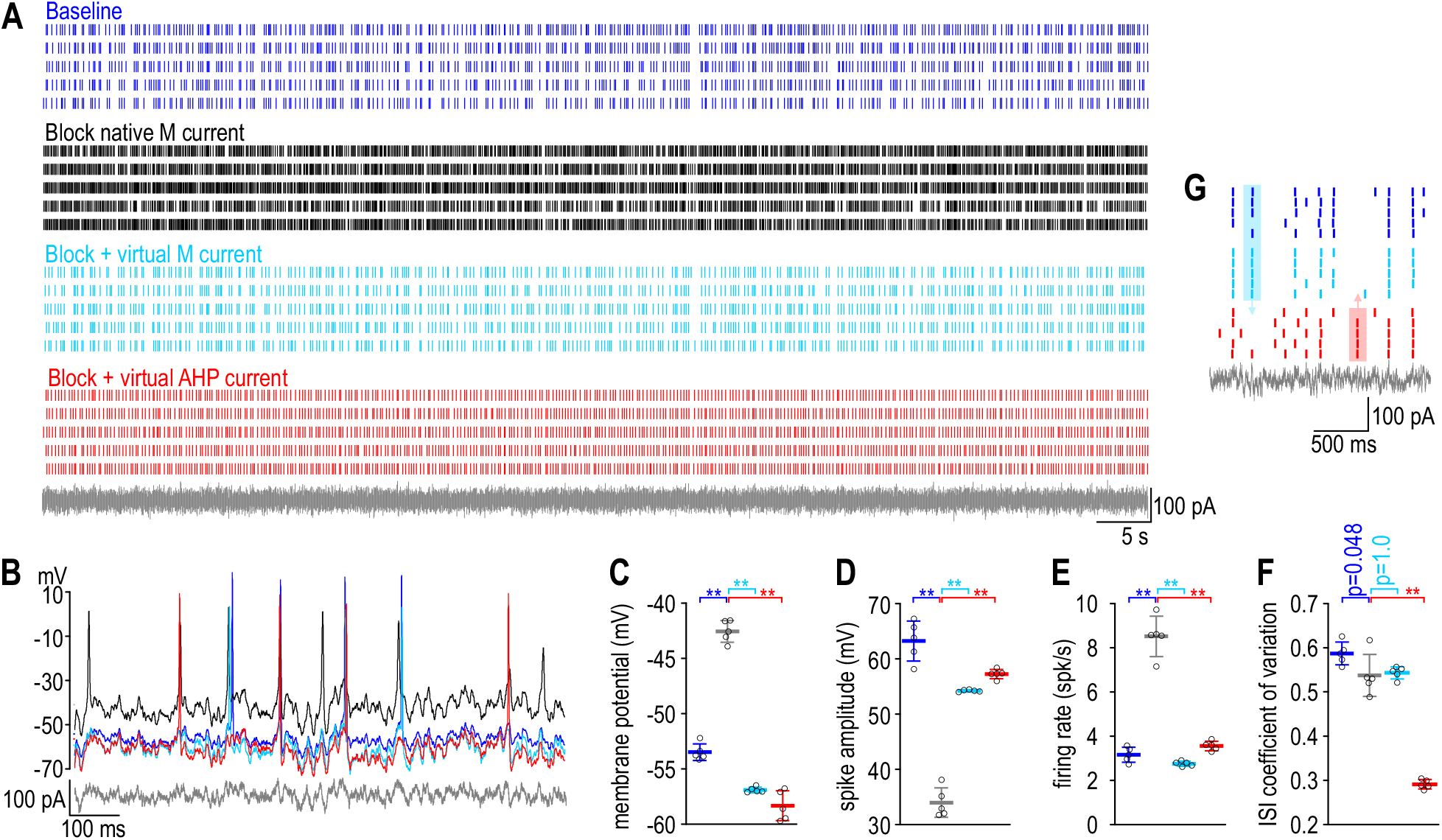
A compensatory change that restores firing rate disrupts firing pattern in a CA1 pyramidal neuron. **(A)** Rasters show spiking in a single CA1 pyramidal neuron given the same noisy current injection (gray) on five trials under each of four conditions: baseline (blue), after blocking native *I*_M_ with 10 μM XE991 (black), and again after inserting virtual *I*_M_ (cyan) or virtual *I*_AHP_ (red) using dynamic clamp. See Methods for virtual conductance parameters. **(B)** Sample voltage traces under each condition. **(C)** Membrane potential differed across conditions (*F*_3,16_ = 298.49, *p* < 0.001, one-way ANOVA); specifically, it was depolarized by blocking *I*_M_ (*t* = 18.71, *p* < 0.001, Tukey test) but that effect was reversed by inserting virtual *I*_M_ (*t* = 24.56, *p* < 0.001) or virtual *I*_AHP_ (*t* = 27.00, *p* < 0.001). **(D)** Spike amplitude also differed across conditions (*F*_3,16_ = 154.04, *p* < 0.001); specifically, it was attenuated by blocking M (*t* = 20.23, *p* < 0.001) but that effect was reversed by inserting virtual *I*_M_ (*t* = 13.99, *p* < 0.001) or virtual *I*_AHP_ (*t* = 16.09, *p* < 0.001). **(E)** Firing rate also differed across conditions (*F*_3,16_ = 177.61, *p* < 0.001); specifically, it was increased by blocking *I*_M_ (*t* = 18.38, *p* < 0.001) but that effect was reversed by inserting virtual *I*_M_ (*t* = 20.85, *p* < 0.001) or virtual *I*_AHP_ (*t* = 16.12, *p* < 0.001). **(F)** Regularity of spiking, reflected in the coefficient of variation of the interspike interval, also differed across conditions (*F*_3,16_ = 188.23, *p* < 0.001), but whereas blocking *I*_M_ had a modest effect (*t* = 2.69, *p* = 0.048) and inserting virtual *I*_M_ had no effect (*t* = 0.39, *p* = 1.00), inserting virtual *I*_AHP_ had a large effect (*t* = 18.23, *p* < 0.001). Data are summarized as mean ± SD. Each data point represents a different trial from a single neuron. **(G)** Enlarged view of rasters to highlight spikes that occurred with native or virtual *I*_M_ but not with virtual *I*_AHP_ (blue shading) or vice versa (red shading) to illustrate the change in spike pattern caused by correcting the change in firing rate caused by blocking *I*_M_ with a compensatory change in *I*_AHP_.

### Few of the ion channel combinations producing the target value for one property also produce the target value for a second property

We proceeded with computational modeling to identify the conditions required to co-regulate multiple properties. For our investigation, it was critical to account for all ion channels that are changing. Even for well characterized neurons like pyramidal cells, there is no comprehensive list of which properties are directly regulated and which ion channel are involved in each case. Therefore, we built a simple model neuron in which >1 property can be co-regulated by adjusting a small number of ion channels. The relative rates at which ion channel densities are updated using the “error” in each property differ between properties, meaning ion channels are co-adjusted according to a certain ratio to regulate one property^8^ but are co-adjusted according to a different ratio to regulate a second property, which has some notable implications (see Discussion). We focused on regulation of independent properties like firing rate and energy efficiency per spike – instead of energy consumption rate, for example, which depends on firing rate – but our conclusions do not hinge on which properties are considered. Target values were chosen arbitrarily.

Simulations were conducted in a single-compartment model neuron whose spikes are generated by a fast sodium conductance and a delayed rectifier potassium conductance with fixed densities. A spike-dependent adaptation mechanism, *g*_AHP_ was included at a fixed density. Densities of all other channels were either systematically varied (and values producing the target output were selected) or they were adjusted via negative feedback to produce the target output (see Methods). The former approach, which amounts to a grid search, identifies all density combinations producing a target output (*i.e.* the solution set). The latter approach finds a subset of those combinations through a homeostatic regulation mechanism. Ion channels with adjustable densities included a sodium conductance (*g*_Na_) and potassium conductance (*g*_K_) that both activate with kinetics similar to the delayed rectifier channel, a slower-activating M-type potassium conductance (*g*_M_), and a leak conductance (*g*_leak_).

We began by testing whether ion channel density combinations that produce the target value for one property also produce a consistent value for other properties. **Figure 3A** shows rheobase for different combinations of 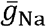 and 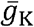. The contour from point 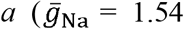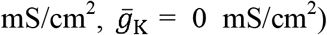 to point 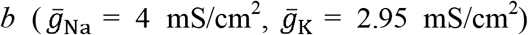 represents all combinations (*i.e.* the solution set) yielding a rheobase of 30 μA/cm^2^. Despite yielding the same rheobase, density combinations along the *a-b* contour did not yield the same minimum sustainable firing rate (*f*_min_) (**Fig. 3B**). The value of *f*_min_ reflects spike initiation dynamics: *f*_min_ ≫0 spk/s is consistent with class 2 excitability^57^ and operation as a coincidence detector^58^, whereas *f*_min_ ≈ 0 spk/s is consistent with class 1 excitability and operation as an integrator, with several consequences (**Fig. 3–figure supplement 1**). After adding *g*_AHP_ to expand the stimulus range over which fluctuation-driven spikes occur, we plotted the firing rate driven by irregularly fluctuating (noisy) *I*_stim_ for different combinations of 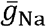 and 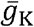 (**Fig. 3C**). Density combinations yielding the same rheobase did not yield equivalent stimulation-evoked firing rates (**Fig. 3D**).

**Figure 3.**
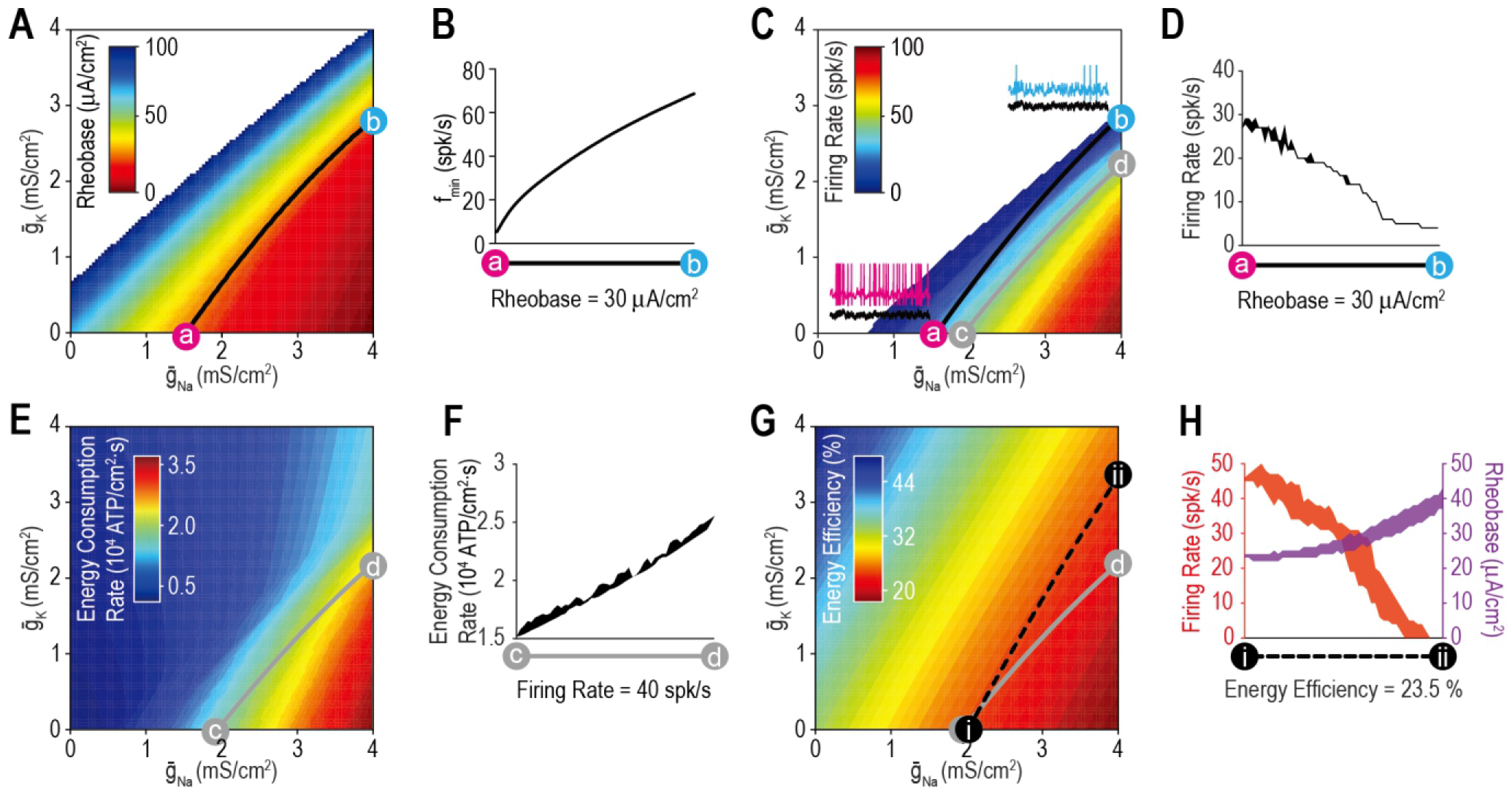
Most ion channel combinations producing the target value for one property produce inconsistent values for other properties. **(A)** Color shows the minimum *I*_stim_ required to evoke repetitive spiking (rheobase) for different combinations of 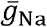 and 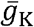. Contour linking *a* and *b* highlights density combinations yielding a rheobase of 30 μA/cm^2^. (B) Minimum sustainable firing rate (*f*_min_) varied along the iso-rheobase contour (see Fig. 3–figure supplement 1). **(C)** Color shows firing rate evoked by noisy *I*_*stim*_ (*τ*_stim_ = 5 ms, *σ*_stim_ = 10 μA/cm^2^, *μ*_stim_= 40 μA/cm^2^) for different combinations of 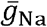 and 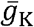. Adding spike-dependent and -independent forms of adaptation 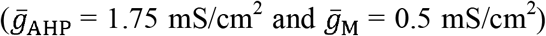 broadened the dynamic range. Grey curve shows density combinations yielding a firing rate of 40 spk/s (*i.e.* an iso-firing rate contour; shown in red in Figs. 4-6). Insets show sample responses to equivalent noisy stimulation. **(D)** Firing rate varied along the iso-rheobase contour from panel A. **(E)** Color shows energy consumption rate for different combinations of 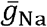 and 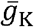 based on firing rates shown in panel C. Iso-firing rate contour *c-d* (grey line in panel C) does not align with energy contours. ATP consumed by the Na^+^/K^+^ pump was calculated from the total Na^+^ influx and K^+^ efflux determined from the corresponding currents. **(F)** Energy consumption rate varied along the iso-firing rate contour. **(G)** Color shows energy efficiency per spike for different combinations of 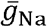 and 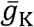. Energy efficiency was calculated as the capacitive minimum, C·ΔV, divided by total Na^+^ influx, where *C* is capacitance and Δ*V* is spike amplitude (see Fig. 3–figure supplement 2). Density combinations along contour *i-ii* (dashed line) yield energy efficiency of 23.5% (shown in green in Fig. 5). **(H)** Both rheobase and firing rate varied along the iso-energy efficiency contour.

Having demonstrated that ion channel density combinations yielding the target value for one aspect of excitability (*e.g.* rheobase) yield differing values for other aspects of excitability (*e.g. f*_min_ or firing rate), we predicted that the same lack of generalization would extend to other cellular properties (*e.g.* energy efficiency). Spikes are energetically costly^59^ but vary in their energy efficiency based on the temporal overlap in sodium and potassium channel activation^60^ (**Fig. 3–figure supplement 2**). Using responses reported in Figure 3C, we measured energy consumption rate for combinations of 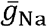 and 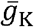 (**Fig. 3E**). Energy consumption rate increased with firing rate, as expected, but did not increase equivalently across all density combinations, as evident from the variation in energy consumption rate along the iso-firing rate contour (**Fig. 3F**). This is due to differences in energy efficiency (**Fig. 3G**) determined as the energy consumed per spike relative to the theoretical minimum (see Methods). Density combinations yielding equally efficient spikes yielded different stimulation-evoked firing rates and rheobase values (**Fig. 3H**). One might presume that spikes are produced as efficiently as possible rather than being regulated to a specific value, but energy efficiency decreases propagation safety factor^61^ and a target value likely emerges by balancing these competing interests^62^.

### Adjusting an ion channel to regulate one property risks disrupting other properties – simulations

Like the experiment in Figure 2, we tested if restoring firing rate to its target value via compensatory changes in either of two ion channels (**Fig. 4A**) after perturbing a third channel would disrupt energy efficiency in our model neuron. Grey circles on **Figure 4B** show randomly chosen combinations of 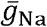 and 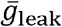 that yield a firing rate of 40 spk/s when 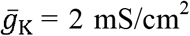. When 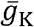 was “blocked” (abruptly reset to 0 mS/cm^2^), firing rate jumped to ~93 spk/s before being restored to 40 spk/s via feedback control of either 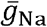 (pink) or 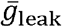 (cyan) (**Fig. 4C**). As 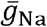 or 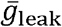 converged on new, compensated densities, firing rate returned to its target value but energy efficiency was affected in opposite ways (**Fig. 4D**). If energy efficiency is also regulated, then 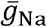 or 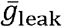 must be co-adjusted in a way that restores firing rate without disrupting energy efficiency.

**Figure 4.**
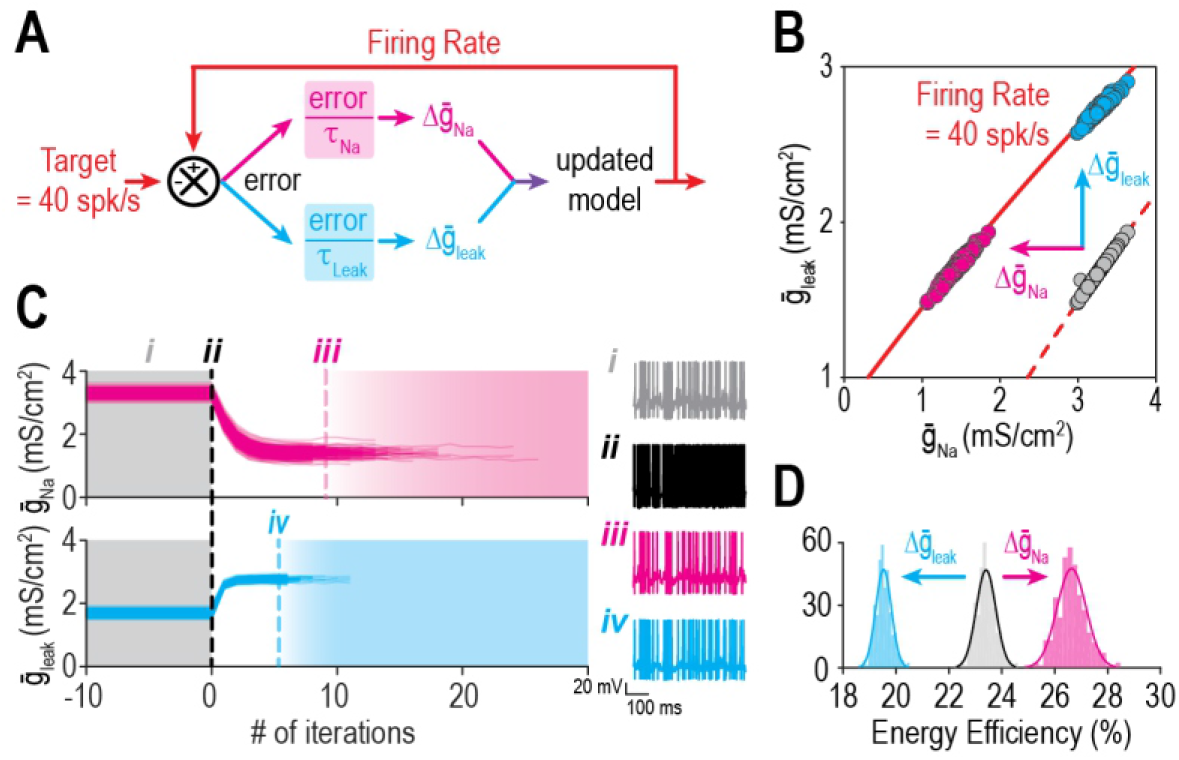
Ion channel changes mediating the same effect on firing rate can oppositely affect energy efficiency. **(A)** Schematic shows how a difference in firing rate from its target value creates an error that is reduced by updating 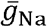 or 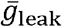. **(B)** Iso-firing rate contours for 40 spk/s are shown for different combinations of 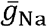 and 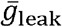 with 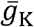 at baseline (2 mS/cm^2^, dashed red curve) and after 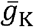 was “knocked out” (0 mS/cm^2^, solid red curve). When 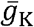 was abruptly reduced, starting models (grey dots) spiked rapidly (~93 spk/s) before firing rate was regulated back to its target value by compensatory changes in either 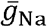 (pink) or 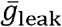 (cyan). Models evolved in different directions and settled at different positions along the solid curve. **(C)** Trajectories show evolution of 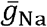 and 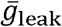. Trajectories are terminated once target firing rate is reached. Sample traces show responses before (grey) and immediately after (black) 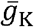 was reduced, and again after compensatory changes in 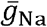 (pink) or 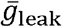 (cyan). **(D)** Distributions of energy efficiency are shown before (grey) and after firing rate regulation via control of 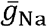 (pink) or 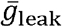 (cyan).

### Geometrical explanation for the relationship between single- and multi-output solutions

Next, we sought a geometrical explanation for how to co-adjust >1 ion channel to co-regulate >1 property. The top panel of **Figure 5A** shows all combinations of 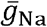 and 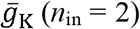 yielding the target value for firing rate (red) *or* energy efficiency (green) (*n*_out_ = 1). Like in Figures 3 and 4, the solution set for each property corresponds to a curve. The multi-output solution set for firing rate *and* energy efficiency (*n*_out_ = 2) corresponds to where the two single-output solution sets intersect, which occurs at a point – only one density combination yields the target values for both properties, meaning the multi-output solution is unique. But if another conductance like *g*_leak_ is adjustable (*n*_in_ = 3), the curves in 2D parameter space (top panel) transform into surfaces in 3D parameter space (bottom panel) and those surfaces intersect along a curve – many density combinations yield the target values for both properties, meaning the multi-output solution is degenerate. The same patterns are evident for firing rate *and* input resistance (**Fig. 5B**) or energy efficiency *and* input resistance (**Fig. 5C**). **Figure 5D** shows that if all three properties – firing rate, energy efficiency *and* input resistance – are regulated (*n*_out_ = 3), the multi-output solution set is empty for *n*_in_ = 2 (*i.e.* the three curves do not intersect at a common point [top panel] unless 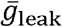 is reset to 1.95 mS/cm^2^ [inset]) and is unique for *n*_in_ = 3 (bottom panel). From this, we conclude that robust regulation of *n* properties requires >*n* adjustable ion channels even if each ion channel contributes to regulation of >1 property. This is like a system of linear equations, which is said to be *underdetermined* if unknowns (inputs) outnumber equations (outputs) (see Discussion).

**Figure 5.**
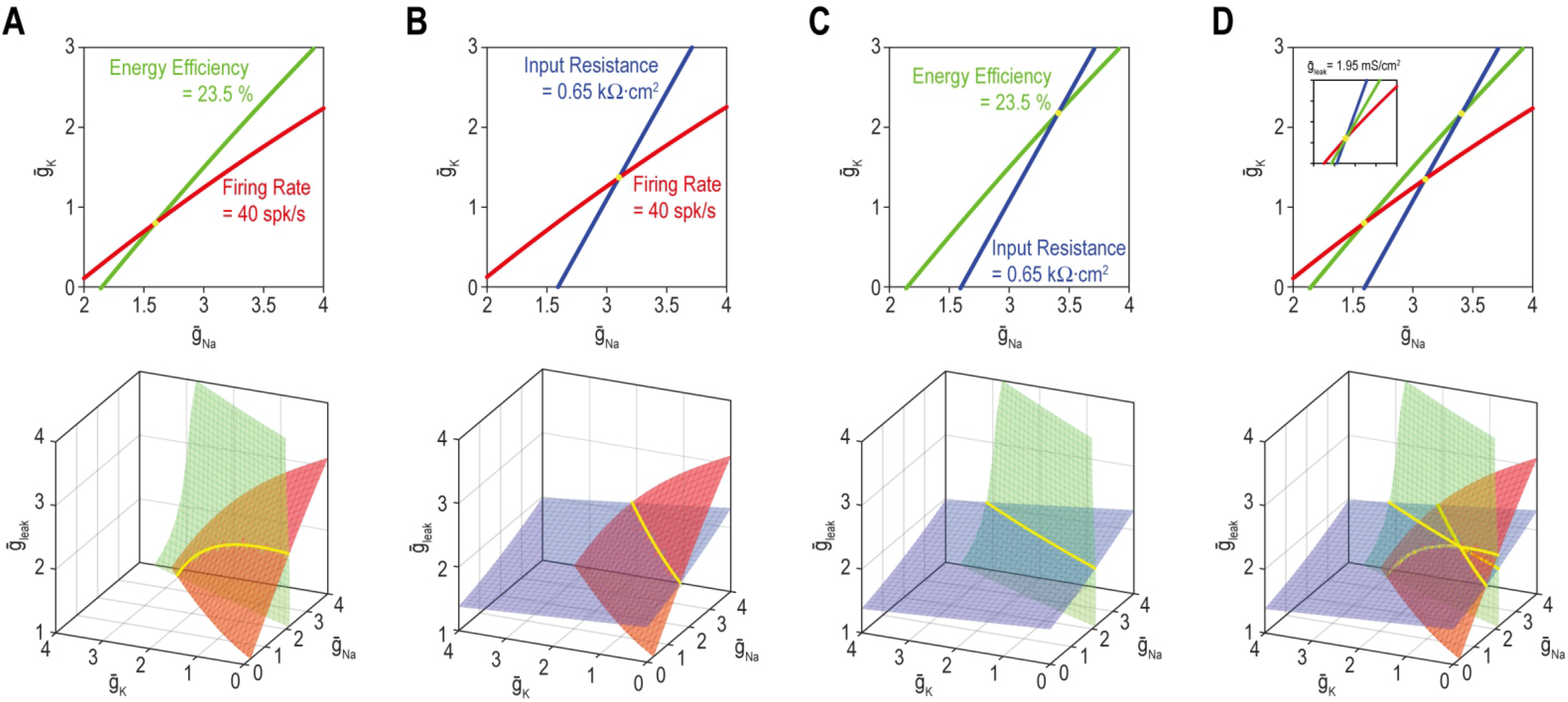
A degenerate solution for *n* properties requires at least *n* + 1 adjustable ion channels. Curves in top panels depict single-output solutions sets based on control of 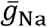 and 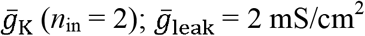. Surfaces in bottom panels depict single-output solution sets based on control of 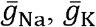, and 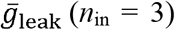. Intersection (yellow) of single-output solutions at a point constitutes a unique multi-output solution, whereas intersection along a curve (or higher-dimensional manifold) constitutes a degenerate multi-output solution. **(A)** Curves for firing rate (40 spk/s) and energy efficiency (23.5 %) intersect at a point whereas the corresponding surfaces intersect along a curve. Solutions for firing rate and input resistance (0.65 kΩ·cm^2^) **(B)** and for energy efficiency and input resistance **(C)** follow the same pattern as in panel A. **(D)** For *n*_in_ = 2 (top), curves for firing rate, energy efficiency and input resistance do not intersect at a common point unless 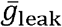 is reset to 1.95 mS/cm^2^ (inset). For *n*_in_ = 3 (bottom), the three surfaces intersect at the same point as in the inset. See Figure 5–figure supplement 1 for the effects of tolerance on solution sets.

In all simulations involving homeostatic regulation, properties were regulated to within certain bounds of the target rather than to a precise value, despite how solution sets are depicted. When tolerances are depicted, single-output solution sets correspond to thin strips (rather than curves) or shallow volumes (rather than surfaces) in 2D and 3D parameter space, respectively (**Fig. 5–figure supplement 1**). The width, depth, etc. are proportional to the tolerance. How does this impact multi-output solutions? *Thin* 2D strips intersect at a small patch (unlike curves intersecting at a point) but that patch in 2D parameter space (Fig. 5–figure supplement 1A) is unlike the long 1D curve formed by *broad* 2D surfaces intersecting in 3D parameter space (see bottom panels of Fig. 5). Likewise, *shallow* 3D volumes intersect at a narrow tube (unlike surfaces intersecting at a curve) but that tube in 3D parameter space (see Fig. 5–figure supplement 1B) is unlike the broad 2D surface formed by *deep* 3D volumes intersecting in 4D parameter space. In other words, increasing tolerance does not increase solution space dimensionality the same way as increasing *n*_in_.

The need for ≥*n* adjustable channels to regulate *n* properties has been shown before11, 50, 56 and may be obvious to some for mathematical reasons, but the relationship has some important implications (*e.g.* for parameter estimation^63^). The implications for homeostatic regulation have not been thoroughly explored, thus prompting the next steps of our study.

### The dimensionality of solution space affects ion channel correlations

We predicted that the dimensionality of solution space – point (0D), curve (1D), surface (2D), volume (3D), etc. – affects ion channel correlations by limiting the degrees of freedom. To explore this, we measured ion channel correlations within single- and multi-output solution sets found through homeostatic regulation. **Figure 6A** shows all combinations of 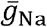 and 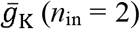 producing the target firing rate (*n*_out_ = 1). Correlation between 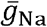 and 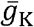 is high because homeostatically determined solutions are constrained to fall along a curve; under those conditions, variation in one channel is offset solely by variation in the other channel, and all covariance is thus captured in a single pairwise relationship. If 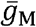 is also allowed to vary (*n*_in_ = 3), homeostatically determined solutions spread across a surface and pairwise correlations become predictably weaker (**Fig. 6B**) since variation in one channel can be offset by variation in two other channels, and covariance is diluted across >1 pairwise relationship. If additional channels are allowed to vary (*n*_in_ ≥ 4), solutions distribute over higher-dimensional manifolds and pairwise correlations further weaken (**Fig. 6C**). However, if firing rate *and* energy efficiency are both regulated (*n*_out_ = 2; **Fig. 6D**), homeostatically determined solutions are once again constrained to fall along a curve when *n*_in_ = 3, and pairwise correlations strengthen (**Fig. 6E**). Increasing *n*_in_ to 4 while keeping *n*_out_ at 2 caused correlations to weaken (Fig. 6F). These results confirm the predicted impact of solution space dimensionality on ion channel correlations.

**Figure 6.**
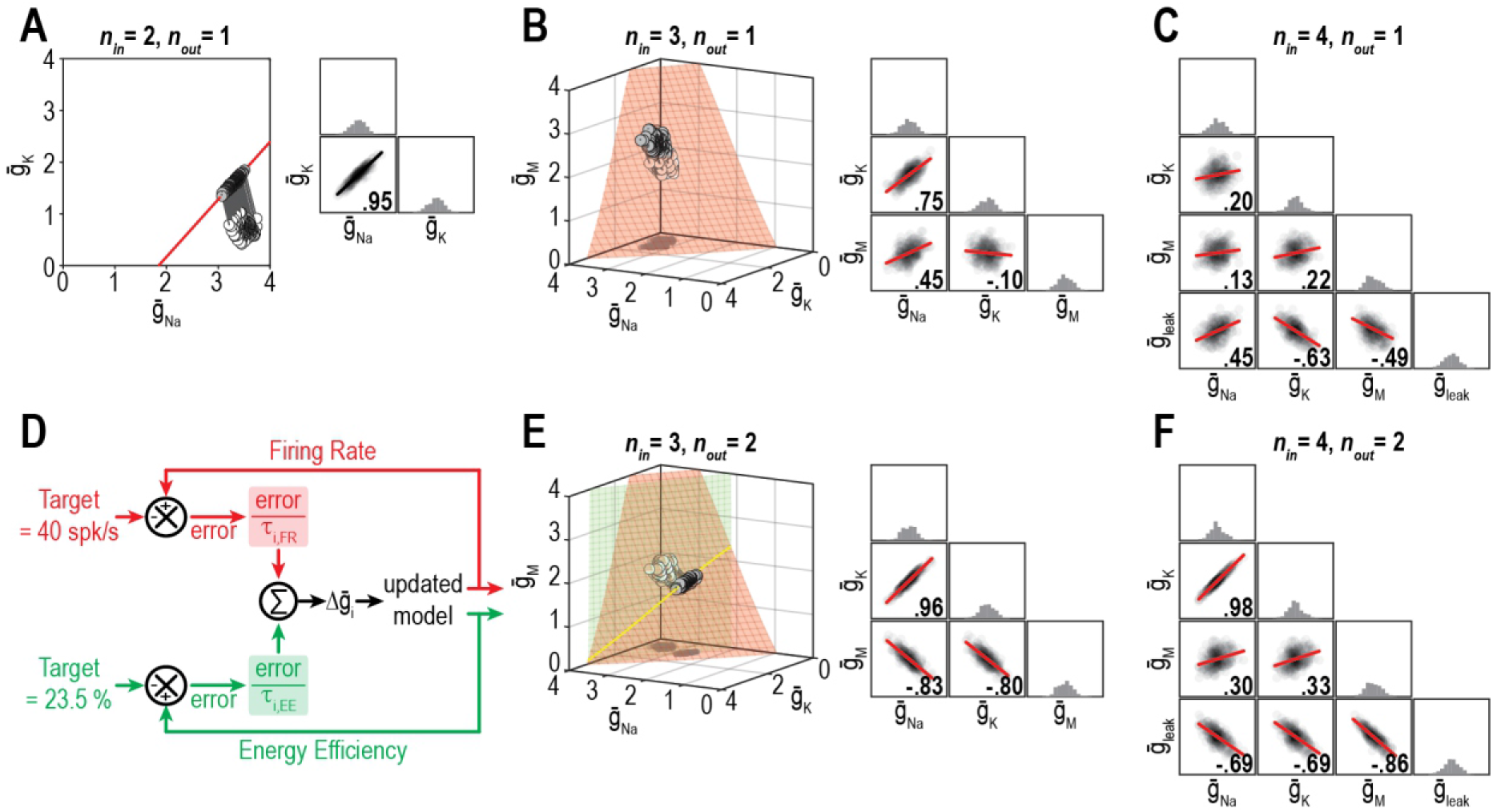
Dimensionality of solution space affects ion channel correlations. **(A)** Starting from a normally distributed cluster of 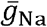 and 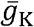 (white dots) yielding an average firing rate of 73 spk/s, 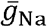 and 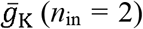 were homeostatically adjusted to regulate firing rate (*n*_out_ = 1) to its target value of 40 spk/s. Grey lines show trajectories. Because solutions (grey dots) converge on a curve, the pairwise correlation between 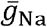 and 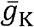 is predictably strong. Scatterplots show solutions centered on the mean and normalized by the standard deviation (z-scores). Correlation coefficient (*R*) is shown in the bottom right corner of each scatter plot. **(B)** Same as panel A (*n*_out_ = 1) but via control of 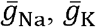 and 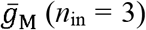. Homeostatically found solutions converge on a surface and ion channel correlations are thus weaker. **(C)** If 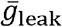 is also controlled (*n*_in_ = 4), solutions converge on a hard-to-visualize volume (not shown) and pairwise correlations are further weakened. **(D)** Schematic shows how errors for two regulated properties are combined: the error for each property is calculated separately and is scaled by its respective control rate *τ* to calculate updates, and all updates for a given ion channel (*i.e.* originating from each error signal) are summed. **(E)** Same as panel B (*n*_in_ = 3), but for regulation of firing rate and energy efficiency (*n*_out_ = 2). Homeostatically found solutions once again converge on a curve (yellow), which now corresponds to the intersection of two surfaces; ion channel correlations are thus strong, like in panel A. **(F)** If *n*_in_ is increased to 4 while *n*_out_ remains at 2, solutions converge on a surface (not shown) and ion channel correlations weaken.

O’Leary et al.^64^ demonstrated how the relative rates at which different ion channel densities are controlled impact their correlation. This is reproduced in **Figure 7A**, where, from the same initial conditions (density combinations), homeostatic control with different relative rates (shown in pink and cyan) produces solutions with different correlations. Relative regulation rates can affect not only the strength of pairwise correlations, but also the sign (compare correlation between 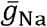 and 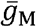). However, if firing rate *and* energy efficiency are both homeostatically regulated, correlations strengthen (consistent with results from Fig. 6) and become independent of the relative regulation rates because solutions are limited to a lower-dimensional solution space (**Fig. 7B**). Recall that dimensionality of the multi-output solution set corresponds to *n*_in_ – *n*_out_. For relative regulation rates to influence ion channel correlations in this way, homeostatically determined solutions must fall on a solution manifold with dimensionality >1, but not ≫1 lest pairwise correlations be diluted.

**Figure 7.**
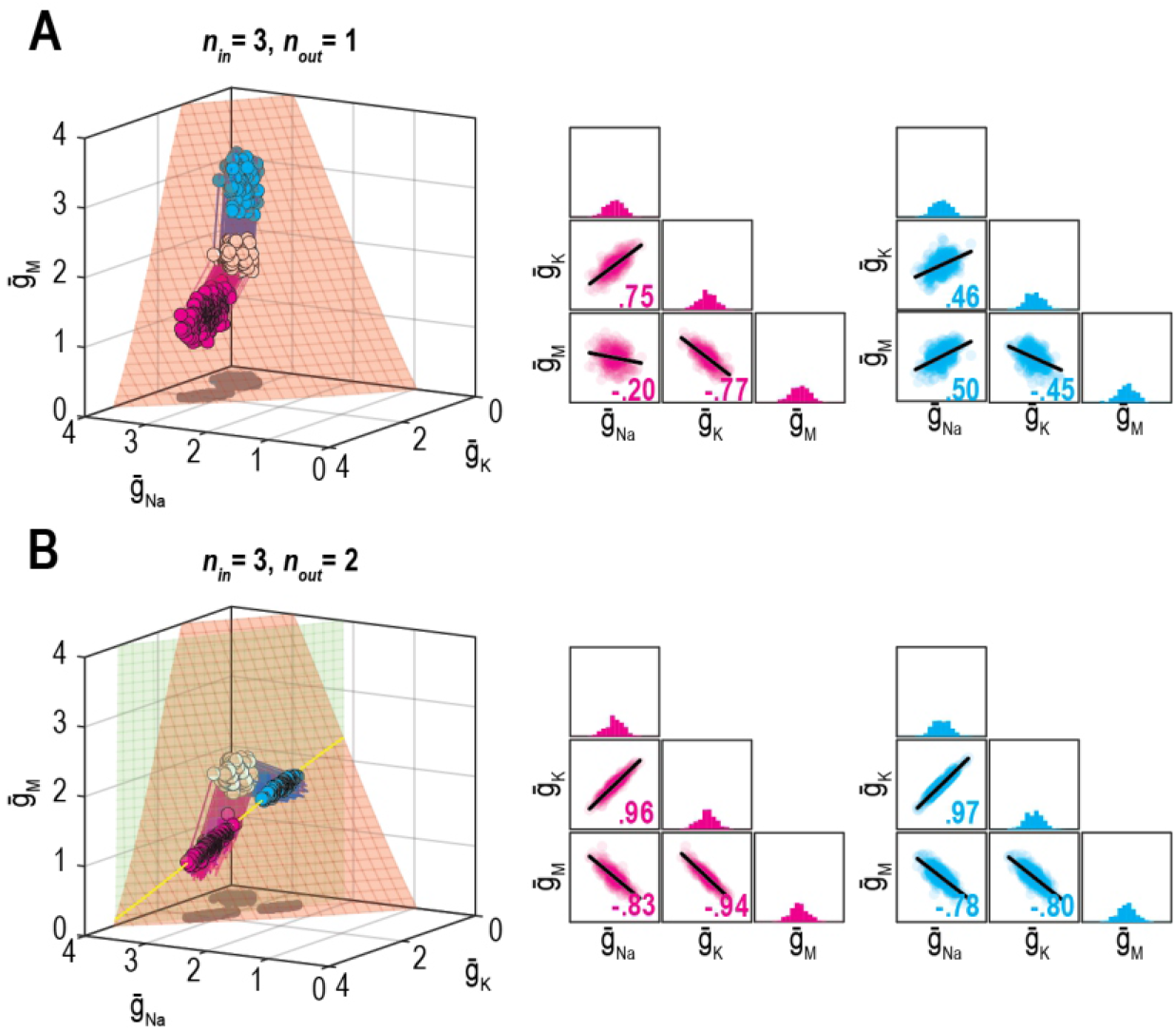
Effect of relative regulation rates on ion channel correlations depends on the dimensionality of solution space. **(A)** Homeostatic regulation of firing rate via control of 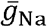, 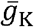 and 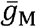. Same as Figure 6B, but for two new sets of regulation rates (see Table S1). Solutions found for each set of rates (pink and cyan) approached the surface from different angles and converged on the surface with different patterns, thus producing distinct ion channel correlations, consistent with O’leary et al.^64^. **(B)** Same as panel A (*n*_in_ = 3), but for homeostatic regulation of firing rate and energy efficiency (*n*_out_ = 2). Solutions converge on a curve (yellow), giving rise to virtually identical ion channel correlations regardless of regulation rates.

However, correlations produced through the homeostatic regulation mechanism described by O’Leary et al.^64^, which is essentially the same mechanism used here, were recently shown to be sensitive to noise^66^. This occurs because this regulation mechanism brings conductance densities to the solution manifold (by minimizing the error signal) but cannot control how solutions drift on the manifold (since the error signal is 0 everywhere on the manifold). Solutions do not drift in the absence of noise but, of course, noise is ubiquitous in biological systems and its effects must be considered. Accordingly, we re-ran simulations from Figures 6 and 7 with noise added to the conductance densities. After a few hundred noise-update iterations, solutions spread across the available solution space, which corresponds to a surface when only firing rate is regulated (**Fig. 8A top**). The distribution of solutions can produce correlations (**Fig. 8A bottom**). Lower and upper bounds on conductance densities shape the solution space and influence correlations (**Fig. 8–figure supplement 1A**). Correlations also depend on how solutions distribute across the solution space, which depends on regulation rates (**Fig. 8–figure supplement 1B,C**), which means correlations can still depend on regulation rates under noisy conditions, but correlations are not the same as in the absence of noise (**Fig. 8B**). When firing rate and energy efficiency were regulated, noise caused solutions to spread across the intersection (**Fig. 8C**) but correlations were relatively unaffected since they are limited by the low-dimensionality of this solution space. Spread was bounded by one or another conductance density reaching 0 mS/cm^2^. Franci et al.^66^ proposed another regulatory scheme in which an attractive subspace emerges through cooperative molecular interactions; correlations created by fluctuations along that subspace are robust to noise. That scheme should preclude correlations from arising as in Figure 8A because solutions are prevented from spreading across the solution space; conversely, a low-dimensional solution space like in Fig. 8C may prevent correlations unless the attractive subspace arising from molecular interactions aligns with the solution space. Further work is required to explore these regulatory mechanisms and how they interact.

**Figure 8.**
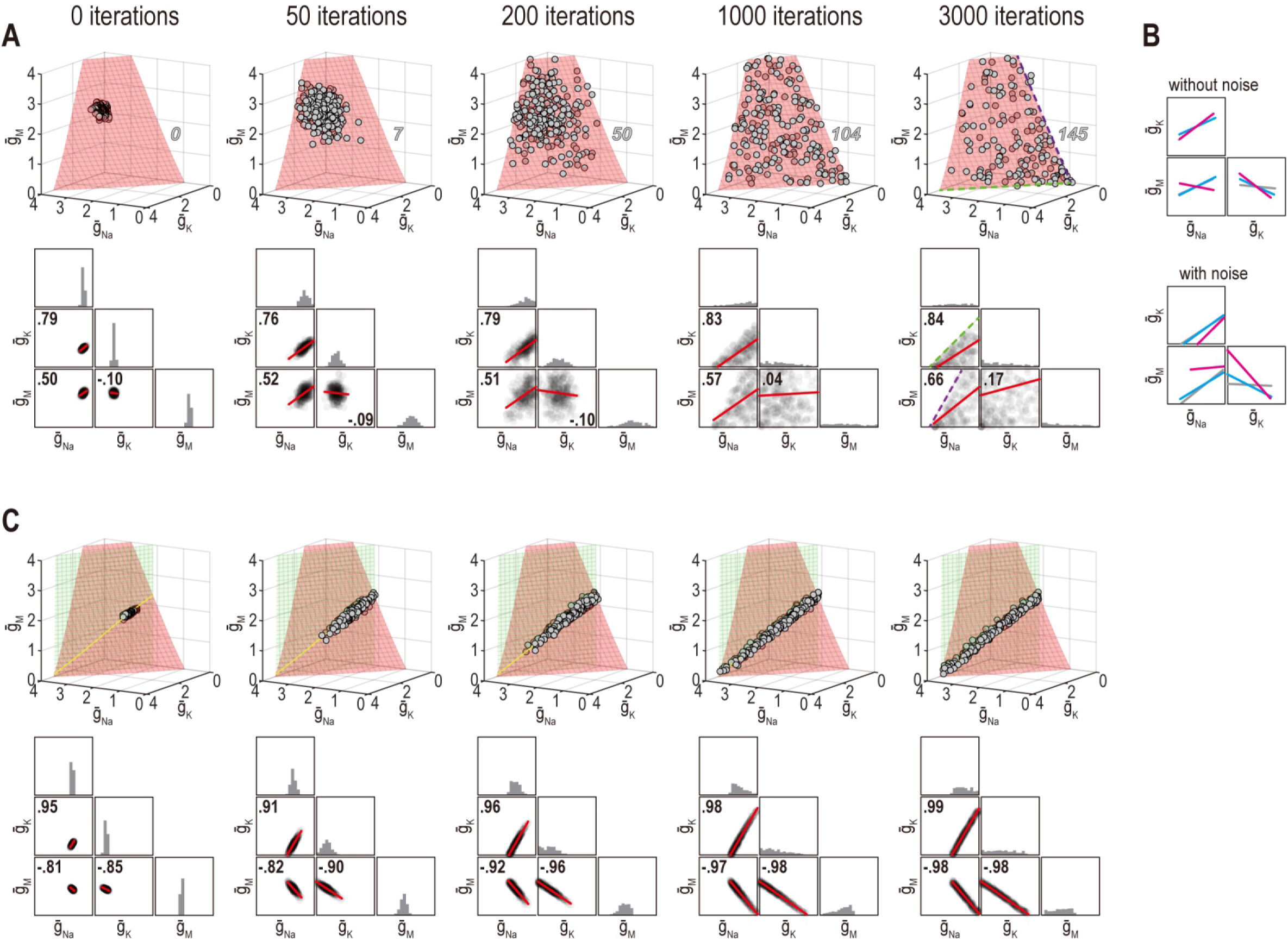
Noise can affect ion channel correlations depending on the dimensionality of solution space. **(A)** Starting from solution sets in Figure 6B, 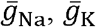 and 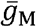 had noise added to them and were then updated by homeostatic regulation (using regulation rates from Figure 6) to correct the noise-induced disruption of firing rate. Conductance density combinations are depicted before and after 50, 200, 1000 or 3000 noise-update iterations (left to right). Noise caused solutions to spread across the surface (top). Some solutions (number in italics) drifted beyond the illustrated region but this is prevented by imposing an upper bound (**Figure 8–figure supplement 1A**). Ion channel correlations reflect the available solution space; for example, changes in 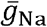 and 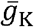 are limited by 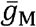 remaining positive (dashed green line), while changes in 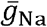 and 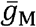 are limited by 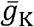 remaining positive (dashed purple line). How solutions distribute within that space depends on regulation rates, and also affects correlations (**Figure 8–figure supplement 1B, C**). Conductance density distributions and pairwise correlations (bottom) were not centered on the mean or normalized by standard deviation (unlike in other figures) in order to visualize how distributions evolve over iterations. **(B)** Comparison of regression lines from Figures 6 (grey) and 7 (pink, cyan) without noise (top) and with noise (bottom). Correlations are affected by regulation rates in both conditions, but not in the same way. **(C)** Same as panel A but for regulation of firing rate *and* energy efficiency. Solutions spread along the intersection of the two surfaces, but the spread is bounded by one or another conductance density reaching 0 mS/cm^2^. Correlations slightly increase under noisy conditions (compare with Figure 6).

### The dimensionality of solution space affects the success of homeostatic regulation

Beyond affecting the ion channel correlations that emerge through homeostatic regulation, we predicted that the dimensionality of solution space affects whether multiple properties can be successfully regulated to their target values. **Figure 9A** shows an example in which co-adjusting 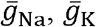, and 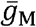 successfully regulates firing rate. **Figure 9B** shows successful regulation of energy efficiency by co-adjusting the same three channels. Using the same initial conditions and relative regulation rates as above, the system failed to co-regulate firing rate *and* energy efficiency (**Fig. 9C**). Notably, coordinated regulation of both properties was achieved using other relative regulation rates (see Figs. 6C and 7B) or using the same relative rates but starting from different initial conditions (not shown), suggesting that low-dimensional solutions are less accessible (*i.e.* a smaller set of relative regulation rates succeed in finding the solution space). This may be a challenge for the Franci et al. regulation model if molecular interactions create an attractive subspace that does not align with a low-dimensional solution space.

**Figure 9.**
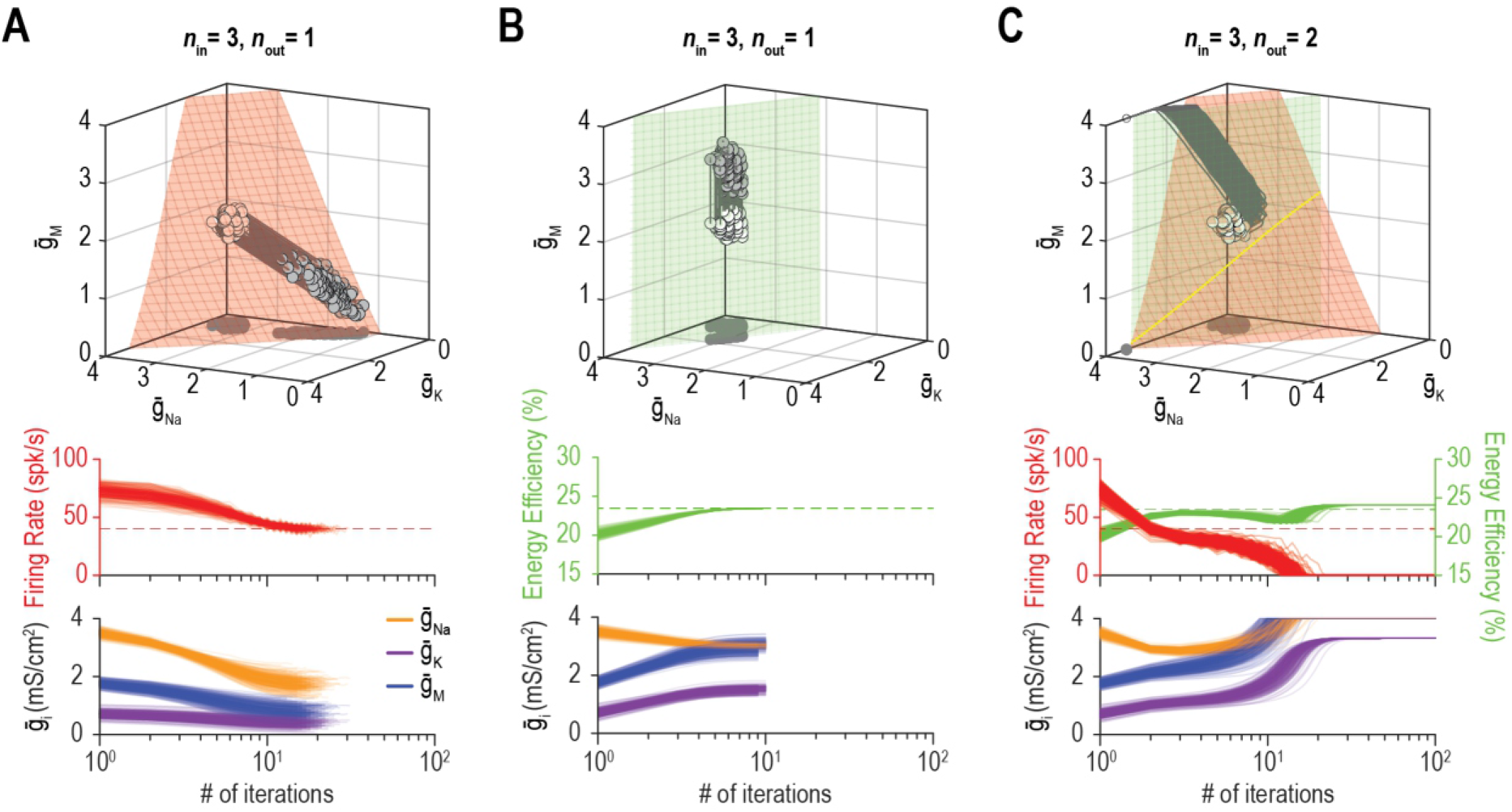
Low-dimensional solutions can be hard for homeostatic regulation to “find”. **(A)** Homeostatic regulation of firing rate (*n*_out_ = 1) via control of 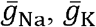 and 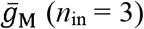, like in Figures 6B and 7A but using a different set of regulation rates (see Table S1). Solutions converged onto the iso-firing rate surface (top panel) and firing rate was regulated to its target value in <30 iterations (bottom panel). **(B)** Same as A (*n*_in_ = 3) but for homeostatic regulation of energy efficiency (*n*_out_ = 1). Solutions converged on the iso-energy efficiency surface (top panel) and energy efficiency was regulated to its target value in ~10 iterations (bottom panel). **(C)** Homeostatic regulation of firing rate and energy efficiency (*n*_out_ = 2) via control of the same ion channels (*n*_in_ = 3) using the same relative rates and initial values as in panels A and B. Neither firing rate (red trajectories) nor energy efficiency (green trajectories) reached its target value. Conductance densities were capped at 4 mS/cm^2^ but this does not account for trajectories not reaching their target. Noise was not included in these simulations.

Figure 10 shows additional examples of regulating firing rate and energy efficiency by co-adjusting 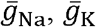, and 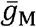. In these simulations, firing rate is regulated to a precise target value whereas energy efficiency is maintained above a lower bound; the single-output solution set for energy efficiency thus corresponds to a volume rather than a surface for *n*_in_ = 3. For energy efficiency ≥22%, homeostatically determined solutions converge on the iso-firing rate surface without regulation of energy efficiency having much effect (**Fig. 10A**). For energy efficiency ≥27%, solutions initially converge onto part of the iso-firing rate surface that sits outside the targeted energy efficiency volume, but without energy efficiency requirements being met, solutions then move across the surface until reaching the intersection with the iso-energy efficiency volume (**Fig. 10B**). By converging on the intersection, which is a curve, ion channel correlations are stronger than in Figure **10**A. In other words, ion channel correlations increase as single-output solution sets start to disconnect, meaning increased correlations presage regulation failure; in general, this is consistent with the impact of increasing constraints (see Fig. 8), which go hand in hand with decreasing degrees of freedom. For energy efficiency ≥30%, the iso-firing rate surface and targeted energy efficiency volume do not intersect and solutions thus proceed to a point between the two single-output solution sets constrained by 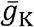 and 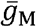 reaching 0 mS/cm^2^ (**Fig. 10C**). In this last example, neither property is optimally regulated but the outcome is a reasonable compromise. Regulation might have failed outright if, in the absence of an upper bound, ion channel densities increased (wound-up) without properties ever reaching their target values. This highlights that co-regulation of multiple properties might fail or find sub-optimal solutions not because single-output solutions do not exist, but because even large single-output solution sets might not intersect, meaning no multi-output solution exists.

**Figure 10.**
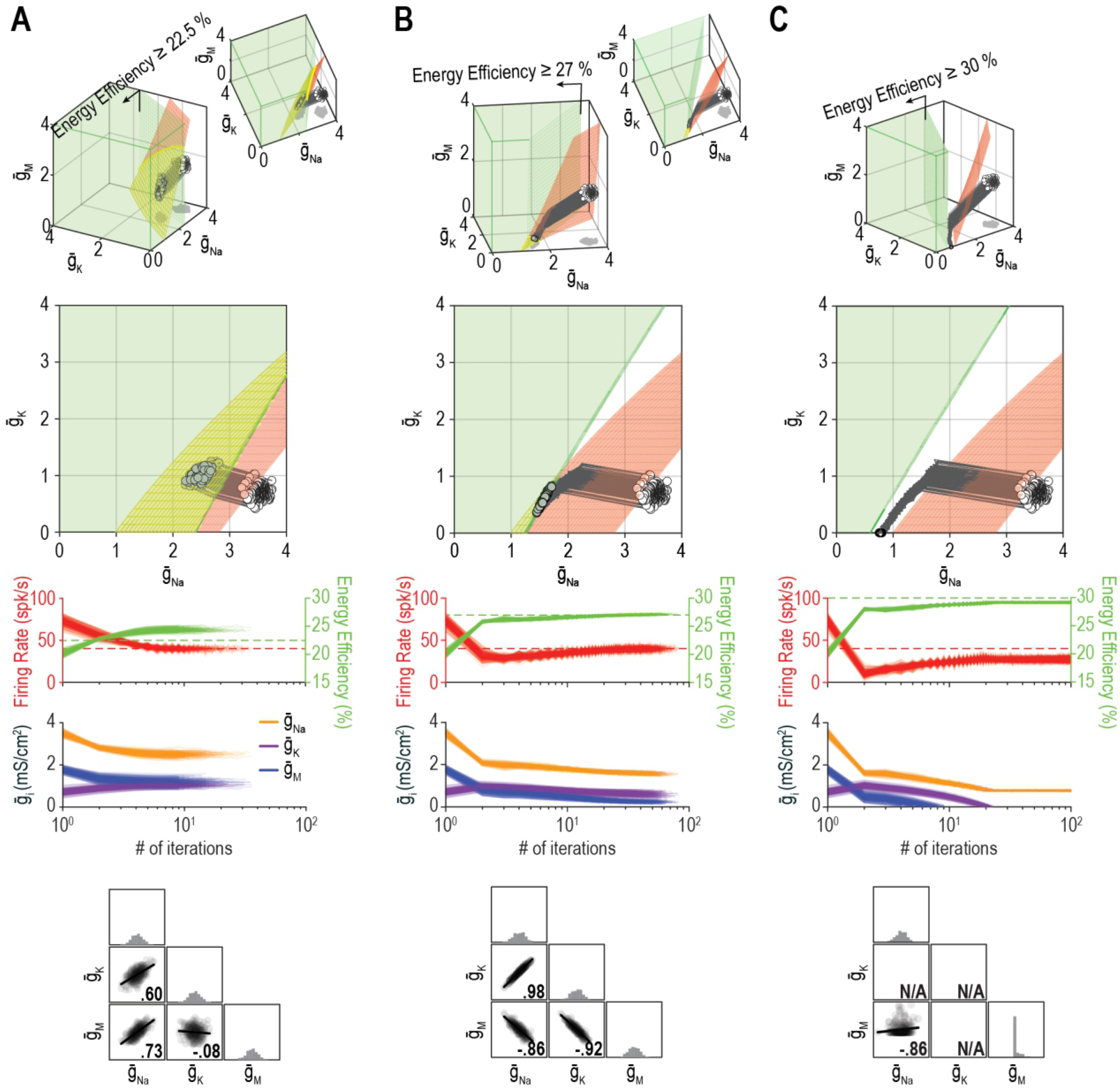
The outcome of homeostatic regulation depends on if and how single-output solution sets intersect. Homeostatic regulation of firing rate and energy efficiency (*n*_out_ = 2) via control of 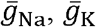 and 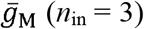. For these simulations, energy efficiency was maintained above a lower bound rather than being regulated to a specific target value; accordingly, the single-output solution set for energy efficiency corresponds to a volume (green) rather than a surface. **(A)** For energy efficiency ≥ 22 %, homeostatically determined solutions converge on the iso-firing rate surface (red) in a region sitting within the green volume (top panel). The rate of convergence and resulting ion channel correlations are shown in the middle and bottom panels, respectively. **(B)** For energy efficiency ≥ 27 %, solutions initially converge on the red surface in a region outside the green volume, but trajectories then bend and proceed across the red surface until that surface reaches the green volume. Because solutions converge on a curve, ion channel correlations are stronger than in A, where solutions distributed across a surface. (C) For energy efficiency ≥ 30 %, the red surface and green volume do not intersect. Consequently, solutions settle between the two single-output solution sets (top panel) without either property reaching its target (middle panel). The outcome represents the balance achieved by the opposing pull of control mechanisms regulating different properties, and depends entirely on 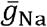 since 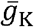 and 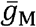 cannot become negative (bottom panel). Noise was not included in these simulations.

## DISCUSSION

This study has identified conditions required for a neuron to co-regulate >1 property. Being able to produce the same output using diverse ion channel combinations ensures that single-output solution sets are large. This is critical because of the many ion channel combinations that produce the desired output for one property, few also produce the desired output for other properties (Fig. 3), which means ion channel adjustments that regulate one property are liable to disrupt other properties due to ion channel pleiotropy (Figs. 2 and 4). Indeed, the multi-output solution set corresponds to the intersection between single-output solution sets and the dimensionality of the multi-output solution corresponds to the difference between the number of adjustable ion channels (*n*_in_) and the number of regulated outputs (*n*_out_) (Fig. 5). Co-regulation of *n* properties requires at least *n* adjustable ion channels for a unique solution, and at least *n*+1 channels for a degenerate solution. This constraint is not alleviated by pleiotropy, but *n*_in_ would need to exceed *n*_out_ by an even wider margin if ion channels were not pleiotropic. Moreover, channels must be co-adjusted with different ratios to regulate different properties, which constrains how feedback loops are organized (see below). These important issues get overlooked if regulation of each property is considered in isolation.

With respect to the number of channels required to co-regulate a certain number of properties, a direct analogy can be made with a system of linear equations. Each unknown constitutes a degree of freedom and each equation constitutes a constraint that reduces the degrees of freedom by one. The system is said to be overdetermined if equations outnumber unknowns, and underdetermined if unknowns outnumber equations. An underdetermined system can have infinite solutions. Degeneracy is synonymous with underdetermination, which highlights that degeneracy depends not only on ion channel diversity, but also on those channels not having to co-regulate “too many” properties. Extra degrees of freedom broaden the range of available solutions, meaning a good solution is liable to exist over a broader range of conditions. On the other hand, if constraints outnumber the degrees of freedom, the system becomes overdetermined and solutions disappear (*e.g.* Fig. 5D), which can cause regulation to fail (see below). One might reasonably speculate that ion channel diversity has been selected for because it enables the addition of new functionality (*e.g.* excitability) without the cell becoming overdetermined, lest pre-existing functions (*e.g.* osmoregulation) become compromised.

There are notable similarities and differences between a neuron adjusting its ion channel densities to regulate properties and a scientist trying to infer conductance densities based on the measured values of those properties (*i.e.* fitting a model to experimental data). Fitting a model with *n* parameters to many outputs (*e.g.* firing rate *and* input resistance *and* rheobase *and* spike height) is more difficult but yields better parameter estimates than fitting the same model to just one output^56^, just as regulating more properties makes the solution set smaller. But how good are those parameter estimates? Assuming there is no measurement noise, can one confidently infer the true conductance densities in a particular neuron by measuring and fitting enough properties? Degeneracy makes solving this inverse problem difficult, if not impossible^63, 65^. The neuron solves the forward problem, producing a firing rate, input resistance, etc. based on its channel density combination. That combination is determined by negative feedback but the precise densities are unimportant so long as the combination produces the target values for all regulated properties (which happens for every density combination in the multi-output solution set). As conditions change, the multi-output solution will evolve and negative feedback will adjust densities accordingly. A neuron needs to regulate its properties but does not do so by regulating its conductance densities to particular values; in that respect, a neuron does not solve an inverse problem. (Convergence of conductance densities to the same “attractive” set regardless of initial conditions, or after a perturbation, may suggest otherwise^66^, but may reflect other, unaccounted for constraints like regulation of another property.) Neurons have evolved under selective pressure to be degenerate (see above), not parsimonious, contrary to how most models are constructed.

Previous studies using grid searches to explore degeneracy have tended to apply selection criteria simultaneously, finding the “multi-output” solution set in one fell swoop. In contrast, we considered one criterion (property) at a time in order to find each single-output solution set, which we then combined to find the multi-output solution. The former approach is akin to aggregating several error functions to create a single-objective problem, whereas the latter resembles multi-objective optimization. Nevertheless, past studies have observed that certain parameter changes (in a particular direction through parameter space) dramatically affect some properties but not others^34, 67^, consistent with manifolds representing the single-output solution sets for sensitive and insensitive properties lying orthogonal to one another in parameter space. The effects of solution dimensionality on ion channel correlations and regulation failure (see below) highlight the value of a multi-objective perspective, which implicitly recognizes that there are multiple properties, each with its own feedback loop, even if those feedback loops intersect. Our results show that co-regulating many properties using a common set of ion channels is feasible only if enough *different* channels are involved. Indeed, for co-adjustments to work, channels must differ from each other in how they affect each property (see Fig. 1), which emphasizes the distinction between degeneracy and redundancy.

Our results also highlight the need for >1 “master regulator” because regulation of each property requires that ion channels are co-adjusted with different ratios (**Fig. 11**). O’Leary et al.^8^ argued in favor of a single master regulator, but that was predicated on using a single factor, namely global calcium, to encode the error signal; because a single factor cannot reach two different targets (*i.e.* simultaneously minimize two errors), it cannot support two master regulators. But local^68^ or kinetically distinct^10^ variations in calcium could encode >1 error signal if the calcium sensors coupling intracellular calcium to gene expression or channel modulation are spatially segregated or operate with different filter properties. Beyond calcium, evidence points to firing rate homeostasis being activity-independent^69^, using membrane potential as feedback^70^, or involving dual feedback loops^16^. Furthermore, the AMP:ATP ratio is used to track energy level^71, 72^ and free amino acids can also be sensed^73^. This is just the tip of the iceberg. Rather than error signals being encoded by a single factor like calcium, we favor the notion of a multi-input/multi-output system^74^ in which factors combine to encode multiple error signals. Take mTOR (mechanistic target of rapamycin) for example; it is modulated by diverse factors and, in turn, modulates many processes including transcription, translation and protein degradation^75^. The regulation model proposed by Franci et al.^66^ provides additional insights. These issues require more investigation but, for now, our results argue that co-regulation of multiple properties is incompatible with the amalgamation of error signals or certain control steps. The analogy with single- and multi-objective optimization (see above) seems apropos. Critically, *n*_in_ exceeding *n*_out_ will not guarantee solutions if degrees of freedom are reduced by bottlenecks elsewhere in the feedback loops.

**Figure 11.**
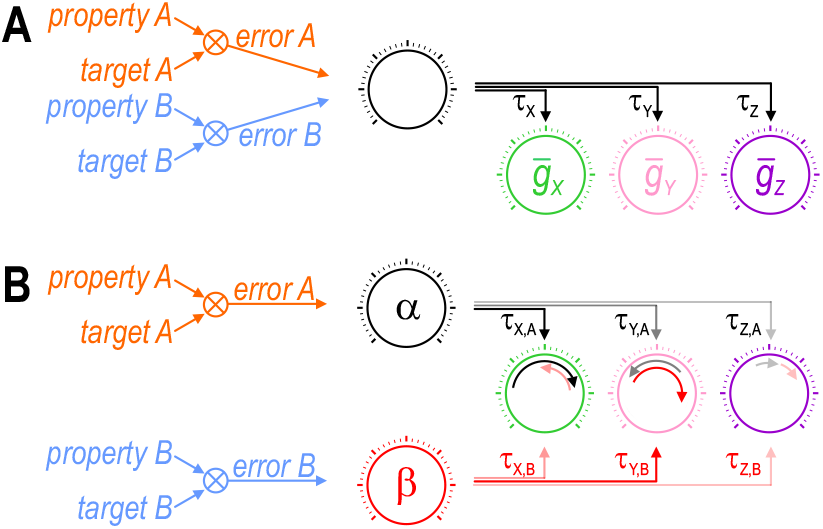
Each error signal must be able to co-adjust channels with different ratios. Cartoons depict regulation of two properties (*A* and *B*) via control of three ion channels (*X*, *Y*, and *Z*). Channel levels are homeostatically adjusted to minimize the difference (error) between each property and its target value. The error signal divided by each channel’s regulation time constant (*τ*) dictates the magnitude of the change in that channel (represented by length of curved arrows in panel B). **(A)** The set of regulation time constants define a “master regulator” (black dial) that co-adjusts ion channels according to a certain ratio. Convergence of error signals at or before the master regulator limits how ion channels are co-adjusted. **(B)** An additional master regulator (red dial) is needed to enable *error B* to co-adjust channels with a different ratio than *error A*. Conditions like in panel B were required for all co-regulation simulations (see Fig. 6C). Different co-adjustment ratios are required to regulate each property because of the different way each ion channel affects each property (see Fig. 1). Note that master regulators may be more distributed than depicted here, with each regulation time constant *τ*_*i,j*_ likely reflecting the net effect of regulating multiple (transcriptional, translational, etc) processes. The positive molecular regulatory network proposed by Franci et al.^66^ involves additional feedback mechanisms not easily depicted in this sort of cartoon.

Ion channel correlations have been studied in simulations^8, 30, 33, 37, 64, 66, 76–78^ and experiments^47, 79–83^. These correlations have been ascribed to the co-regulation of ion channels. The relative rates with which different conductance densities are controlled dictate the direction in which trajectories move through parameter space, which in turn dictates how trajectories approach and distribute across the solution manifold^64^. Our results are consistent with that explanation (Fig. 7), notwithstanding noise effects (see below), but they also highlight the importance of manifold dimensionality: Correlations are stronger but are insensitive to relative regulation rates if trajectories reach a 1D manifold (curve) rather than a higher dimensional manifold (surface, volume, etc.) (Fig. 6). Under noisy conditions, correlations reflect the available solution space and how solutions drift across it, which depends on regulation rates (Fig. 8). The cooperative molecular interactions proposed by Franci et al.^66^ can produce correlations in a high-dimensional solution space, but may be prevented from doing so by a low-dimensional solution space. Notwithstanding molecular interactions, the existence of pairwise correlations suggests that the dimensionality of solution space (= *n*_in_ – *n*_out_) is relatively low. This may be surprising since *n*_in_ is high, but makes sense if *n*_out_ is also high; in other words, there are *many* channels but they are responsible for regulating *many* properties. Though *n*_out_ is constrained by *n*_in_ (*i.e.* the number of regulated properties cannot exceed the number of adjustable ion channels), neurons might take advantage of regulating as many properties as their ion channel diversity safely allows, leading to a relatively low dimensional solution space (as *n*_out_ approaches *n*_in_).

Homeostatic regulation can fail for different reasons. If there are no solutions (*i.e.* the solution set is empty), negative feedback cannot regulate a property to its target values. A multi-output solution set can be empty because single-output solution sets do not intersect (Fig. 10C), meaning regulation of different properties to their respective target values is incompatible. Regulation can also fail because negative feedback fails to converge on available solutions (Fig. 9C). Notably, a multi-output solution set may become less accessible (lower dimensional) as single-output solution sets start to separate (Fig. 10B), and may foreshadow the eventual disjunction of those solution sets (Fig. 10C). Regulation can also fail because feedback signaling is compromised. These failure modes are not mutually exclusive; for example, a system might reach less accessible solutions if regulation rates are normally flexible but might fail to reach those solutions if regulation rate flexibility is reduced. Cooperative molecular interactions^66^ are notable in this regard, insofar as they would tend to constrain regulation and thus influence failure modes. Failed homeostatic regulation may have similar consequence regardless of how the failure occurs, consistent with the emergence of common disease phenotypes despite vastly different underlying pathologies^84^, but subtle differences in exactly how the failure transpires may provide important clues to help pinpoint the underlying mechanism(s).

In conclusion, neurons can co-regulate multiple properties by co-adjusting ion channels. Despite reusing channels to regulate more than one property, many channels are required to co-regulate many properties and feedback loops must allow for co-adjustments with different ratios. This may account for why evolution has yielded such diverse channels and why their transcriptional regulation has become so complicated^85^. Given how difficult it is to study regulation of any one property, studying the co-regulation of multiple properties seems truly daunting, especially if feedback loops intermingle and error signals involve combinatorial codes. But if that is indeed the case, then accounting for the co-regulation of multiple properties may be crucial for appreciating design features that might otherwise elude us.

## METHODS

### Slice electrophysiology

All procedures were approved by the Hospital for Sick Children Animal Care Committee. Using the same procedures and equipment previously described^86^, coronal slices of hippocampus were prepared from an adult mouse and a CA1 pyramidal neuron was recorded using whole-cell patch clamp. A junction potential correction of −9 mV was applied to the recorded membrane potential. A noisy stimulus was generated through an Ornstein-Uhlenbeck process (see Eqn 7) with *τ*_stim_ = 5 ms, *μ*_stim_ = 60 pA, and *σ*_stim_ = 10 pA; the same stimulus was re-played on each trial. Native *I*_M_ was blocked by bath application of 10 uM XE991 (Tocris). The block was continued throughout dynamic clamp experiments. Virtual *I*_M_ and *I*_AHP_ were modeled as per Eqn 6 below (with *τ*_z_ = 100 ms, *β*_z_ = −35 mV and *γ*_z_ = 4 mV for *I*_M_, and *τ*_z_ = 600 ms, *β*_z_ = 0 mV and *γ*_z_ = 1 mV for *I*_AHP_) and were applied using the dynamic clamp capabilities of Signal v6 (Cambridge Electronic Design). The density of each virtual conductance was adjusted to produce the desired firing rates 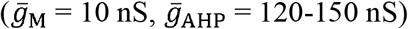.

### Neuron model

Our base model includes a fast-activating sodium conductance (*g*_fast_) and a slower-activating potassium conductance (*g*_slow_) which together are sufficient to produce spikes. Other conductances were added to modulate excitability.

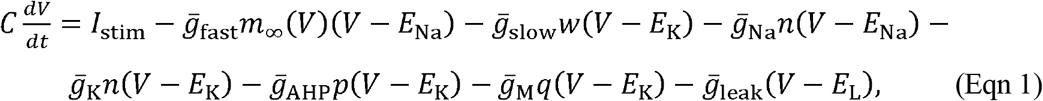

where *V* is voltage and *m* changes instantaneously with *V* whereas other gating variables change more slowly. The spike-generating conductances *g*_*fast*_ and *g*_*slow*_ were modeled using a Morris-Lecar formalism^45^, with *C* = 2 µF/cm^2^, *E*_Na_= 50 mV, *E*_K_= −100 mV, *E*_leak_ = −70 mV, *φ*_w_= 0.15, 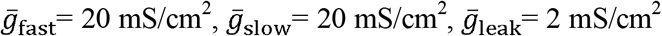, *β*_m_= −1.2 mV, *γ*_m_= 18 mV, *β*_w_ = −10 mV, and *γ*_w_= 10 mV. A generic sodium conductance (*g*_Na_) and potassium conductance (*g*_K_) were modeled using Hodgkin-Huxley formalism as described by Ratté et al.^35^

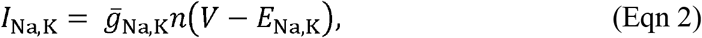

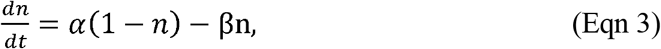

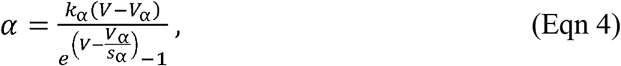

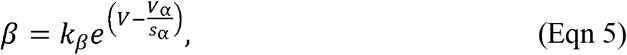

where *V*_α, β_ = −24 mV, *s*_α,β_ = −17 mV, and *k*_α,β_ = 1 ms^−1^. Note that *g*_Na_ and *g*_K_ differ only in their reversal potentials. Channels approximating a calcium-activated potassium conductance (*g*_AHP_) and an M-type potassium conductance (*g*_M_) were modeled as described by Prescott et al.^87^

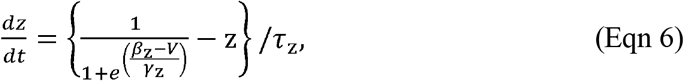

where *τ*_z_= 100 ms, *γ*_z_ = 4 mV, and *β*_z_ = 0 mV or −35 mV for *g*_AHP_ and *g*_M_, respectively. Maximal conductance densities for *g*_leak_, *g*_Na_, *g*_K_ and *g*_M_ were systematically varied or adjusted by a homeostatic feedback mechanism (see below). Any other parameters that differ from the sources cited above are reported in the relevant figure legends. Injected current *I*_stim_ was applied as either a constant step or as noisy fluctuations modeled with an Ornstein-Uhlenbeck process

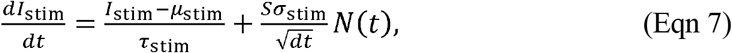

where *τ*_stim_ is a time constant that controls the rate at which *I*_stim_ drifts back towards the mean *μ*_stim_, and *N*(*t*) is a random number drawn from a normal distribution with 0 mean and unit variance that is scaled by *Sσ*_stim_, where a scaling factor 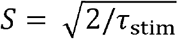 makes the standard deviation *σ*_stim_ independent of *τ*_stim_. All simulations were conducted in MATLAB using the forward Euler integration method and a time step of 0.05-0.1 ms.

### Energy calculations

Energy consumption rate was calculated as in Hasenstaub et al.^88^ Briefly, models were stimulated with a fast fluctuating stimulus (*µ*_stim_ = 40 µA/cm^2^, *σ*_stim_ = 10 μA/cm^2^). Sodium and potassium current through all channels was integrated for 1 second to determine the charge for each ion species, which was then converted to ion flux based on the elementary charge 1.602 × 10^−19^ C. Based on the 3:2 stoichiometry of the Na^+^/K^+^ pump, we divided the number of sodium and potassium ions by 3 and 2, respectively, and used the maximum of those two values as the energy consumption rate (in ATP/cm^2^·s). Energy efficiency was calculated as the ratio between capacitive minimum and total Na^+^ flux during an action potential^60^. Briefly, voltage was reset to −40 mV to evoke a single spike in all models. Models were treated as pure capacitors to calculate the minimum capacitive current as *C*Δ*V*, where *C* is the capacitance and Δ*V* is the difference between the resting membrane potential and the spike peak.

### Grid search

Models were tested with conductance density combinations chosen from a 100×100 or 30×30×30 grid for 2D and 3D plots, respectively. To depict each single-output solution set, all models with outputs within a range (tolerance) of the target value (± 3 spk/s for firing rate, ±0.25 % for energy efficiency, and ± 0.003 kΩ·cm^2^ for input resistance) were selected and a curve or surface was fit to those successful models. The same tolerances were implemented in the homeostatic learning rule (see below) to minimize ringing. Tolerances are illustrated in Figure 5 – figure supplement 1 but were not shown in other figures for sake of clarity.

### Feedback control

We used a homeostatic learning rule similar to O’Leary et al.^8, 64^ with two notable differences: (1) we did not use intracellular calcium or other biological signals (e.g. AMP:ATP ratio) as intermediaries for our error signals and (2) the error for each output was determined as the difference between the current and target values at the end of each iteration, and conductance densities were adjusted before the start of the next iteration, rather than updating and feeding back error signals in “real time”. The separation between fast and slow (regulatory) timescales justifies the former approach. For each conductance 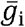, error was divided by the regulation time constant (τ_i_) and added to the conductance (see Fig. 4A). For co-regulating >1 property, we used the sum of scaled errors to update conductance densities (see Fig. 6D). A single run consists of a maximum of 200 iterations, during which a model must reach and maintain its regulated property within the tolerance for five consecutive iterations. All models that reached the target output(s) did so in well under 100 iterations; regulation was deemed to have failed for models not reaching their target output(s) within 200 iterations. Conductance densties during the last five iterations were averaged and reported as the final value. See **Table S1** for the initial conductance densities and regulation time constants used for each figure.

### Conductance noise

For simulations in Figure 8, noisy variations in each conductance density, 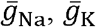 and 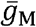, was applied by adding a random number drawn from a Gaussian distribution with a mean of 0 and a standard deviation of 0.05 mS/cm^2^. Noise was independent for each conductance. After applying noise, the simulation was run, errors were calculated, and conductance densities were updated according to the feedback control described above; this qualifies as one noise-update iteration. Up to 3000 noise-update iterations were run. If the addition of noise caused a conductance density to become negative, the density was reset to 0 mS/cm^2^ before applying feedback control. Where indicated, conductance density was likewise reset to 4 mS/cm^2^ if noise caused it to increase above that value.

## CODE AVAILABILITY

All code will be made available at GitHub and ModelDB upon publication.

## ACKNOWLEDGEMENTS

This research was funded by a Discovery Grant from the Natural Sciences and Engineering Research Council (NSERC) and by a Foundation Grant from the Canadian Institutes of Health Research (CIHR). H.S. was supported by Ontario Graduate Scholarship, the University of Toronto Centre for the Study of Pain Scholarship, and James F. Crothers Family Scholarship. J.Y. was supported by a SickKids Restracomp Studentship and Ontario Graduate Scholarship. We thank Eve Marder, Etay Hay, Shreejoy Tripathy, and Simon Hardy for constructive feedback on the manuscript.

## AUTHOR CONTRIBUTIONS

J.Y., H.S. and S.A.P. conceived and designed the study. J.Y. and H.S. conducted simulations S.R. conducted experiments. J.Y., H.S. and S.R. analyzed data. J.Y. and S.A.P. wrote the final paper with feedback from H.S and S.R.

## FIGURE LEGENDS

**Figure 3–figure supplement 1. Ion channel combinations yielding equivalent rheobase produce different responses to noisy stimulation, reflecting fundamental differences in spike initiation dynamics and operating model. (A)** Firing rate is shown as a function of *I*_stim_ for combination *a* (pink), *b* (cyan), and other values tested at 0.5 mS/cm^2^ increments in 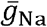 stimulation modeled as an Ornstein-Uhlenbeck (OU) process with stim = 5 ms and astim = 5, 1 (black) along the iso-rheobase contour *a* - *b* in Figure 3. Grey arrows point to responses to noisy stimulation modeled as an Ornstein-Uhlenbeck (OU) process with *τ*_stim_ = 5 ms and (*σ*_stim_ = 5, 1 or 2 μA/cm^2^ and *μ*stim = 25, 29.5 or 35 μA/cm^2^ for sub-, peri- and suprathreshold conditions (panels B - D, respectively). **(B)** Spike-triggered averages (STAs) show that model *a* operates as an integrator with a longer integration time than model *b*, which operates as a coincidence model *b*, consistent with differences in rmin, and average membrane potential differed by >10 detector. **(C)** Noise-induced membrane potential oscillations were slower for model *a* than for model *b*, consistent with differences in *f*_min_, and average membrane potential differed by >10 mV. **(D)** Spike-dependent adaptation 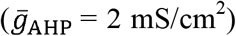 caused bursting in model *b* but not in model *a*. Voltage threshold also differed by 10 mV.

**Figure 3–figure supplement 2. Overlap between Na^+^ and K^+^ currents dictates energy efficiency.** Voltage, total Na^+^ current and total K^+^ current during an action potential are shown for the most efficient (left) and least efficient (right) models. Simultaneous activation of Na^+^ and K^+^ channels creates “waste” current because Na^+^ influx and K^+^ efflux cancel each other out. Overlap between Na^+^ and K^+^ currents is larger in the model on the right, meaning more current is used relative to the theoretical minimum required to spike, which is calculated as *C*·Δ*V*, where *C* is the membrane capacitance and Δ*V* is the difference between the resting membrane potential and the spike peak.

**Figure 5–figure supplement 1. Increasing tolerance does not increase the dimensionality of multi-output solution sets the same way as increasing *n*_in_. (A)** Curves for firing rate (40 spk/s) and energy efficiency (23.5 %) expand into strips when tolerance (±3 spk/s and ± 0.25 %, respectively) is depicted. The strips intersect as a patch (yellow highlighting), unlike curves which intersect at a point (see Fig. 5A, top). Note that 2D strips are limited to a 2D parameter space. The broad 2D surfaces that exist in 3D parameter space intersect along a long curve (see Fig. 5A, bottom), unlike the small patch. **(B)** If 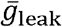 is also controllable (*n*_in_ = 3), strips in 2D parameter space expand into shallow volumes in 3D parameter space. Those shallow volumes intersect along a narrow tube (yellow highlighting), which is unlike the broad surface formed by the intersection of deep volumes in 4D parameter space (not illustrated).

**Figure 8–figure supplement 1. Correlations depend on the solution manifold’s shape and how solutions distribute across it. (A)** Same conditions as in Figure 8A but with conductance densities restricted from increasing above 4 mS/cm^2^, thus preventing solutions from drifting beyond the illustrated region. An upper bound on conductance densities almost certainly exist since transcript and protein levels reflect an equilibrium of production and degradation, where rate-limiting production steps saturate^89^. Imposing an upper bound further constrains the spread of solutions and shapes correlations; for example, changes in 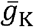 and 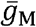 are constrained by 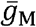 remaining below 4 mS/cm^2^ (dashed blue line). **(B)** Effect of noise after 3000 iterations without (left) and with (right) an upper bound, using cyan regulation rates from Figure 7. **(C)** Same as right panel in B but using pink regulation rates from Figure 7. Noise causes solutions to spread across the manifold but solutions can drift preferentially in a certain direction depending on regulation rates; for example, solutions drift upward and yield high values of 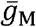 for pink regulation rates, whereas the opposite occurred for the other regulation rates. The distribution of points affects correlations (see Fig. 8B).

## REFERNCES

1. Turrigiano, G., Abbott, L.F. & Marder, E. Activity-dependent changes in the intrinsic properties of cultured neurons. Science 264, 974–977 (1994).

2. O’Leary, T., van Rossum, M.C. & Wyllie, D.J. Homeostasis of intrinsic excitability in hippocampal neurones: dynamics and mechanism of the response to chronic depolarization. J Physiol 588, 157–170 (2010).

3. Aizenman, C.D., Akerman, C.J., Jensen, K.R. & Cline, H.T. Visually driven regulation of intrinsic neuronal excitability improves stimulus detection in vivo. Neuron 39, 831–842 (2003).

4. Desai, N.S., Rutherford, L.C. & Turrigiano, G.G. Plasticity in the intrinsic excitability of cortical pyramidal neurons. Nat. Neurosci. 2, 515–520 (1999).

5. Hengen, K.B., Lambo, M.E., Van Hooser, S.D., Katz, D.B. & Turrigiano, G.G. Firing rate homeostasis in visual cortex of freely behaving rodents. Neuron 80, 335–342 (2013).

6. van, W.I., van Hooft, J.A. & Wadman, W.J. Background activity regulates excitability of rat hippocampal CA1 pyramidal neurons by adaptation of a K+ conductance. J Neurophysiol 95, 2007–2012 (2006).

7. Joseph, A. & Turrigiano, G.G. All for one but not one for all: Excitatory synaptic scaling and intrinsic excitability are coregulated by CaMKIV, whereas inhibitory synaptic scaling is under independent control. J Neurosci 37, 6778–6785 (2017).

8. O’Leary, T., Williams, A.H., Franci, A. & Marder, E. Cell types, network homeostasis, and pathological compensation from a biologically plausible ion channel expression model. Neuron 82, 809–821 (2014).

9. O’Leary, T. & Marder, E. Temperature-Robust Neural Function from Activity-Dependent Ion Channel Regulation. Curr Biol 26, 2935–2941 (2016).

10. Liu, Z., Golowasch, J., Marder, E. & Abbott, L.F. A model neuron with activity-dependent conductances regulated by multiple calcium sensors. J Neurosci 18, 2309–2320 (1998).

11. Olypher, A.V. & Prinz, A.A. Geometry and dynamics of activity-dependent homeostatic regulation in neurons. J Comput Neurosci 28, 361–374 (2010).

12. LeMasson, G., Marder, E. & Abbott, L.F. Activity-dependent regulation of conductances in model neurons. Science 259, 1915–1917 (1993).

13. O’Leary, T. & Wyllie, D.J. Neuronal homeostasis: time for a change? J Physiol 589, 4811–4826 (2011).

14. Davis, G.W. Homeostatic control of neural activity: from phenomenology to molecular design. Annu Rev Neurosci 29, 307–323 (2006).

15. Turrigiano, G. Too many cooks? Intrinsic and synaptic homeostatic mechanisms in cortical circuit refinement. Annu Rev Neurosci 34, 89–103 (2011).

16. Kulik, Y., Jones, R., Moughamian, A.J., Whippen, J. & Davis, G.W. Dual separable feedback systems govern firing rate homeostasis. Elife 8, e45717 (2019).

17. Maffei, A. & Turrigiano, G.G. Multiple modes of network homeostasis in visual cortical layer 2/3. J Neurosci 28, 4377–4384 (2008).

18. Marder, E. Variability, compensation, and modulation in neurons and circuits. Proc Natl Acad Sci U S A 108 Suppl 3, 15542–15548 (2011).

19. Edelman, G.M. & Gally, J.A. Degeneracy and complexity in biological systems. Proc Natl Acad Sci U S A 98, 13763–13768 (2001).

20. Marder, E. & Goaillard, J.M. Variability, compensation and homeostasis in neuron and network function. Nat Rev Neurosci 7, 563–574 (2006).

21. Cropper, E.C., Dacks, A.M. & Weiss, K.R. Consequences of degeneracy in network function. Curr Opin Neurobiol 41, 62–67 (2016).

22. O’Leary, T. Homeostasis, failure of homeostasis and degenerate ion channel regulation. Curr Opin Physiol 2, 129–138 (2018).

23. Ratté, S. & Prescott, S.A. Afferent hyperexcitability in neuropathic pain and the inconvenient truth about its degeneracy. Curr Opin Neurobiol 36, 31–37 (2016).

24. Tononi, G., Sporns, O. & Edelman, G.M. Measures of degeneracy and redundancy in biological networks. Proc Natl Acad Sci U S A 96, 3257–3262 (1999).

25. Goaillard, J.M. & Dufour, M.A. The pros and cons of degeneracy. Elife 3, e02615 (2014).

26. Mason, P.H., Dominguez, D.J., Winter, B. & Grignolio, A. Hidden in plain view: degeneracy in complex systems. Biosystems 128, 1–8 (2015).

27. Rathour, R.K. & Narayanan, R. Degeneracy in hippocampal physiology and plasticity. Hippocampus 29, 980–1022 (2019).

28. Klassen, T., et al. Exome sequencing of ion channel genes reveals complex profiles confounding personal risk assessment in epilepsy. Cell 145, 1036–1048 (2011).

29. Trojanowski, N.F., Padovan-Merhar, O., Raizen, D.M. & Fang-Yen, C. Neural and genetic degeneracy underlies Caenorhabditis elegans feeding behavior. J Neurophysiol 112, 951–961 (2014).

30. Mukunda, C.L. & Narayanan, R. Degeneracy in the regulation of short-term plasticity and synaptic filtering by presynaptic mechanisms. J Physiol 595, 2611–2637 (2016).

31. Anirudhan, A. & Narayanan, R. Analogous synaptic plasticity profiles emerge from disparate channel combinations. J Neurosci 35, 4691–4705 (2015).

32. Kim, E.Z., Vienne, J., Rosbash, M. & Griffith, L.C. Nonreciprocal homeostatic compensation in Drosophila potassium channel mutants. J Neurophysiol 117, 2125–2136 (2017).

33. Taylor, A.L., Goaillard, J.M. & Marder, E. How multiple conductances determine electrophysiological properties in a multicompartment model. J Neurosci 29, 5573–5586 (2009).

34. Drion, G., O’Leary, T. & Marder, E. Ion channel degeneracy enables robust and tunable neuronal firing rates. Proc Natl Acad Sci U S A 112, E5361–5370 (2015).

35. Ratté, S., Zhu, Y., Lee, K.Y. & Prescott, S.A. Criticality and degeneracy in injury-induced changes in primary afferent excitability and the implications for neuropathic pain. Elife 3, 02370 (2014).

36. Mittal, D. & Narayanan, R. Degeneracy in the robust expression of spectral selectivity, subthreshold oscillations, and intrinsic excitability of entorhinal stellate cells. J Neurophysiol 120, 576–600 (2018).

37. Jain, A. & Narayanan, R. Degeneracy in the emergence of spike-triggered average of hippocampal pyramidal neurons. Scientific reports 10, 374 (2020).

38. Gunay, C., Edgerton, J.R. & Jaeger, D. Channel density distributions explain spiking variability in the globus pallidus: a combined physiology and computer simulation database approach. J Neurosci 28, 7476–7491 (2008).

39. Migliore, R., et al. The physiological variability of channel density in hippocampal CA1 pyramidal cells and interneurons explored using a unified data-driven modeling workflow. PLoS Comput Biol 14, e1006423 (2018).

40. Grashow, R., Brookings, T. & Marder, E. Compensation for variable intrinsic neuronal excitability by circuit-synaptic interactions. J Neurosci 30, 9145–9156 (2010).

41. Marder, E. & Taylor, A.L. Multiple models to capture the variability in biological neurons and networks. Nat Neurosci 14, 133–138 (2011).

42. Prinz, A.A., Bucher, D. & Marder, E. Similar network activity from disparate circuit parameters. Nat Neurosci 7, 1345–1352 (2004).

43. Price, C.J. & Friston, K.J. Degeneracy and cognitive anatomy. Trends Cog Sci 6, 416–421 (2002).

44. Onasch, S. & Gjorgjieva, J. Circuit stability to perturbations reveals hidden variability in the balance of intrinsic and synaptic conductances. J Neurosci 40, 3186–3202 (2020).

45. Rho, Y.A. & Prescott, S.A. Identification of molecular pathologies sufficient to cause neuropathic excitability in primary somatosensory afferents using dynamical systems theory. PLoS Comput Biol 8, e1002524 (2012).

46. Swensen, A.M. & Bean, B.P. Robustness of burst firing in dissociated purkinje neurons with acute or long-term reductions in sodium conductance. J Neurosci 25, 3509–3520 (2005).

47. Zhao, S. & Golowasch, J. Ionic current correlations underlie the global tuning of large numbers of neuronal activity attributes. J Neurosci 32, 13380–13388 (2012).

48. Achard, P. & De Schutter, E. Complex parameter landscape for a complex neuron model. PLoS Comput Biol 2, e94 (2006).

49. Bhalla, U.S. & Bower, J.M. Exploring parameter space in detailed single neuron models: simulations of the mitral and granule cells of the olfactory bulb. J Neurophysiol 69, 1948–1965 (1993).

50. Olypher, A.V. & Calabrese, R.L. Using constraints on neuronal activity to reveal compensatory changes in neuronal parameters. J Neurophysiol 98, 3749–3758 (2007).

51. Stemmler, M. & Koch, C. How voltage-dependent conductances can adapt to maximize the information encoded by neuronal firing rate. Nat Neurosci 2, 521–527 (1999).

52. Frere, S. & Slutsky, I. Alzheimer’s disease: from firing instability to homeostasis network collapse. Neuron 97, 32–58 (2018).

53. Fisher, S.K., Heacock, A.M., Keep, R.F. & Foster, D.J. Receptor regulation of osmolyte homeostasis in neural cells. J Physiol 588, 3355–3364 (2010).

54. Balch, W.E., Morimoto, R.I., Dillin, A. & Kelly, J.W. Adapting proteostasis for disease intervention. Science 319, 916–919 (2008).

55. Goaillard, J.M. & Marder, E. Ion channel degeneracy, variability, and covariation in neuron and circuit resilience. Annu Rev Neurosci (2021).

56. Foster, W.R., Ungar, L.H. & Schwaber, J.S. Significance of conductances in Hodgkin-Huxley models. J Neurophysiol 70, 2502–2518 (1993).

57. Hodgkin, A.L. The local electric changes associated with repetitive action in a non-medullated axon. J Physiol 107, 165–181 (1948).

58. Ratté, S., Lankarany, M., Rho, Y.A., Patterson, A. & Prescott, S.A. Subthreshold membrane currents confer distinct tuning properties that enable neurons to encode the integral or derivative of their input. Front Cell Neurosci 8, 452 (2014).

59. Attwell, D. & Laughlin, S.B. An energy budget for signaling in the grey matter of the brain. J Cereb Blood Flow Metab 21, 1133–1145 (2001).

60. Sengupta, B., Stemmler, M., Laughlin, S.B. & Niven, J.E. Action potential energy efficiency varies among neuron types in vertebrates and invertebrates. PLoS Comput Biol 6, e1000840 (2010).

61. Al-Basha, D. & Prescott, S.A. Intermittent failure of spike propagation in primary afferent neurons during tactile stimulation. J Neurosci 39, 9927–9939 (2019).

62. Speakman, J.R., et al. Set points, settling points and some alternative models: theoretical options to understand how genes and environments combine to regulate body adiposity. Dis Mod Mech 4, 733–745 (2011).

63. Sarkar, A.X. & Sobie, E.A. Regression analysis for constraining free parameters in electrophysiological models of cardiac cells. PLoS Comput Biol 6, e1000914 (2010).

64. O’Leary, T., Williams, A.H., Caplan, J.S. & Marder, E. Correlations in ion channel expression emerge from homeostatic tuning rules. Proc Natl Acad Sci U S A 110, E2645–2654 (2013).

65. Aster, R.C., Borchers, B. & Thurber, C.H. Parameter estimation and inverse problems (Academic Press, Waltham, MA, 2013).

66. Franci, A., O’Leary, T. & Golowasch, J. Positive dynamical networks in neuronal regulation: How tunable variability coexists with robustness. IEEE Control Syst Let 4, 946–951 (2020).

67. Goldman, M.S., Golowasch, J., Marder, E. & Abbott, L.F. Global structure, robustness, and modulation of neuronal models. J Neurosci 21, 5229–5238 (2001).

68. Bootman, M.D., Lipp, P. & Berridge, M.J. The organisation and functions of local Ca2+ signals. J Cell Sci 114, 2213–2222 (2001).

69. MacLean, J.N., Zhang, Y., Johnson, B.R. & Harris-Warrick, R.M. Activity-independent homeostasis in rhythmically active neurons. Neuron 37, 109–120 (2003).

70. Santin, J.M. & Schulz, D.J. Membrane Voltage Is a Direct Feedback Signal That Influences Correlated Ion Channel Expression in Neurons. Curr Biol 29, 1683–1688.e1682 (2019).

71. Hardie, D.G. AMP-activated protein kinase: maintaining energy homeostasis at the cellular and whole-body levels. Annu Rev Nutr 34, 31–55 (2014).

72. Garcia, D. & Shaw, R.J. AMPK: mechanisms of cellular energy sensing and restoration of metabolic balance. Molec Cell 66, 789–800 (2017).

73. Efeyan, A., Zoncu, R. & Sabatini, D.M. Amino acids and mTORC1: from lysosomes to disease. Trends Molec Med 18, 524–533 (2012).

74. Papin, J.A. & Palsson, B.O. Topological analysis of mass-balanced signaling networks: a framework to obtain network properties including crosstalk. J Theor Biol 227, 283–297 (2004).

75. Switon, K., Kotulska, K., Janusz-Kaminska, A., Zmorzynska, J. & Jaworski, J. Molecular neurobiology of mTOR. Neuroscience 341, 112–153 (2017).

76. Soofi, W., Archila, S. & Prinz, A.A. Co-variation of ionic conductances supports phase maintenance in stomatogastric neurons. J Comput Neurosci 33, 77–95 (2012).

77. Hudson, A.E. & Prinz, A.A. Conductance ratios and cellular identity. PLoS Comput Biol 6, e1000838 (2010).

78. Ball, J.M., Franklin, C.C., Tobin, A.E., Schulz, D.J. & Nair, S.S. Coregulation of ion channel conductances preserves output in a computational model of a crustacean cardiac motor neuron. J Neurosci 30, 8637–8649 (2010).

79. Temporal, S., Lett, K.M. & Schulz, D.J. Activity-dependent feedback regulates correlated ion channel mRNA levels in single identified motor neurons. Curr Biol 24, 1899–1904 (2014).

80. Schulz, D.J., Goaillard, J.M. & Marder, E. Variable channel expression in identified single and electrically coupled neurons in different animals. Nat Neurosci 9, 356–362 (2006).

81. Tobin, A.E., Cruz-Bermúdez, N.D., Marder, E. & Schulz, D.J. Correlations in ion channel mRNA in rhythmically active neurons. PLoS One 4, e6742 (2009).

82. Schulz, D.J., Goaillard, J.M. & Marder, E.E. Quantitative expression profiling of identified neurons reveals cell-specific constraints on highly variable levels of gene expression. Proc Natl Acad Sci U S A 104, 13187–13191 (2007).

83. Khorkova, O. & Golowasch, J. Neuromodulators, not activity, control coordinated expression of ionic currents. J Neurosci 27, 8709–8718 (2007).

84. Ramocki, M.B. & Zoghbi, H.Y. Failure of neuronal homeostasis results in common neuropsychiatric phenotypes. Nature 455, 912–918 (2008).

85. Gabel, H.W., et al. Disruption of DNA-methylation-dependent long gene repression in Rett syndrome. Nature 522, 89–93 (2015).

86. Khubieh, A., Ratté, S., Lankarany, M. & Prescott, S.A. Regulation of cortical dynamic range by background synaptic noise and feedforward inhibition. Cereb Cortex 26, 3357–3369 (2016).

87. Prescott, S.A. & Sejnowski, T.J. Spike-rate coding and spike-time coding are affected oppositely by different adaptation mechanisms. J Neurosci 28, 13649–13661 (2008).

88. Hasenstaub, A., Otte, S., Callaway, E. & Sejnowski, T.J. Metabolic cost as a unifying principle governing neuronal biophysics. Proc Natl Acad Sci U S A 107, 12329–12334 (2010).

89. Schwanhäusser, B., et al. Global quantification of mammalian gene expression control. Nature 473, 337–342 (2011).

