## Supplemental figures and table for "Minimal requirements for a neuron to co-regulate many properties and the implications for ion channel correlations and robustness"

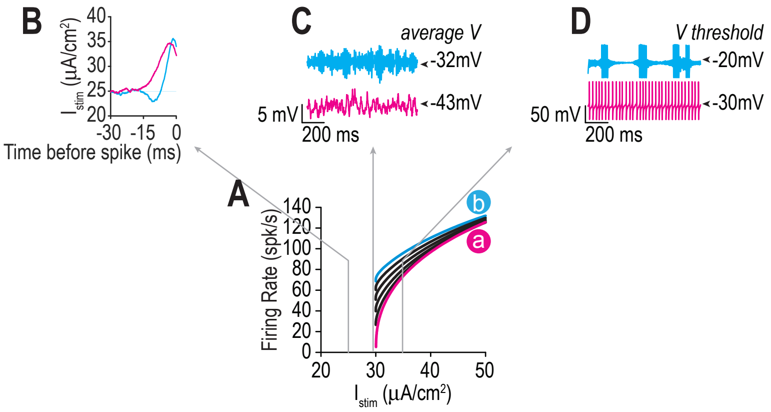


**Figure 3 - figure supplement 1**

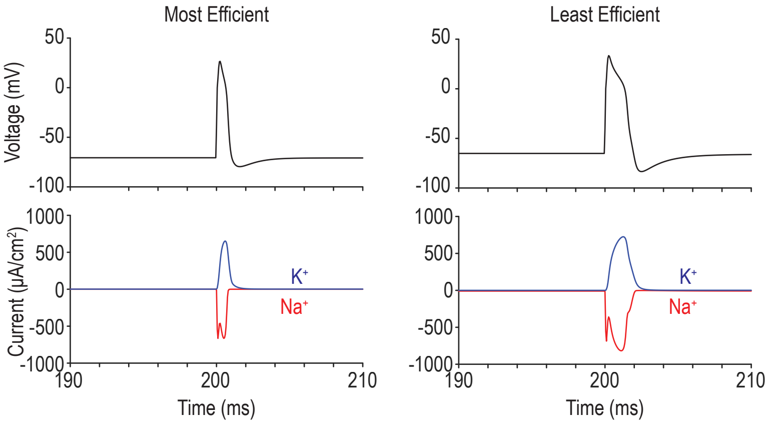


**Figure 3 - figure supplement 2**

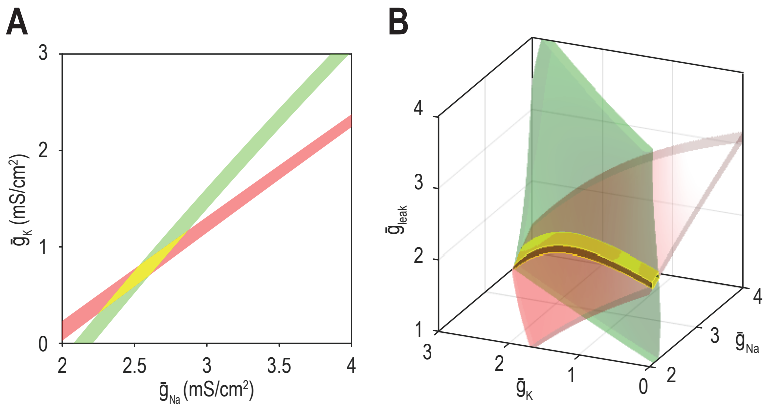


**Figure 5 - figure supplement 1**

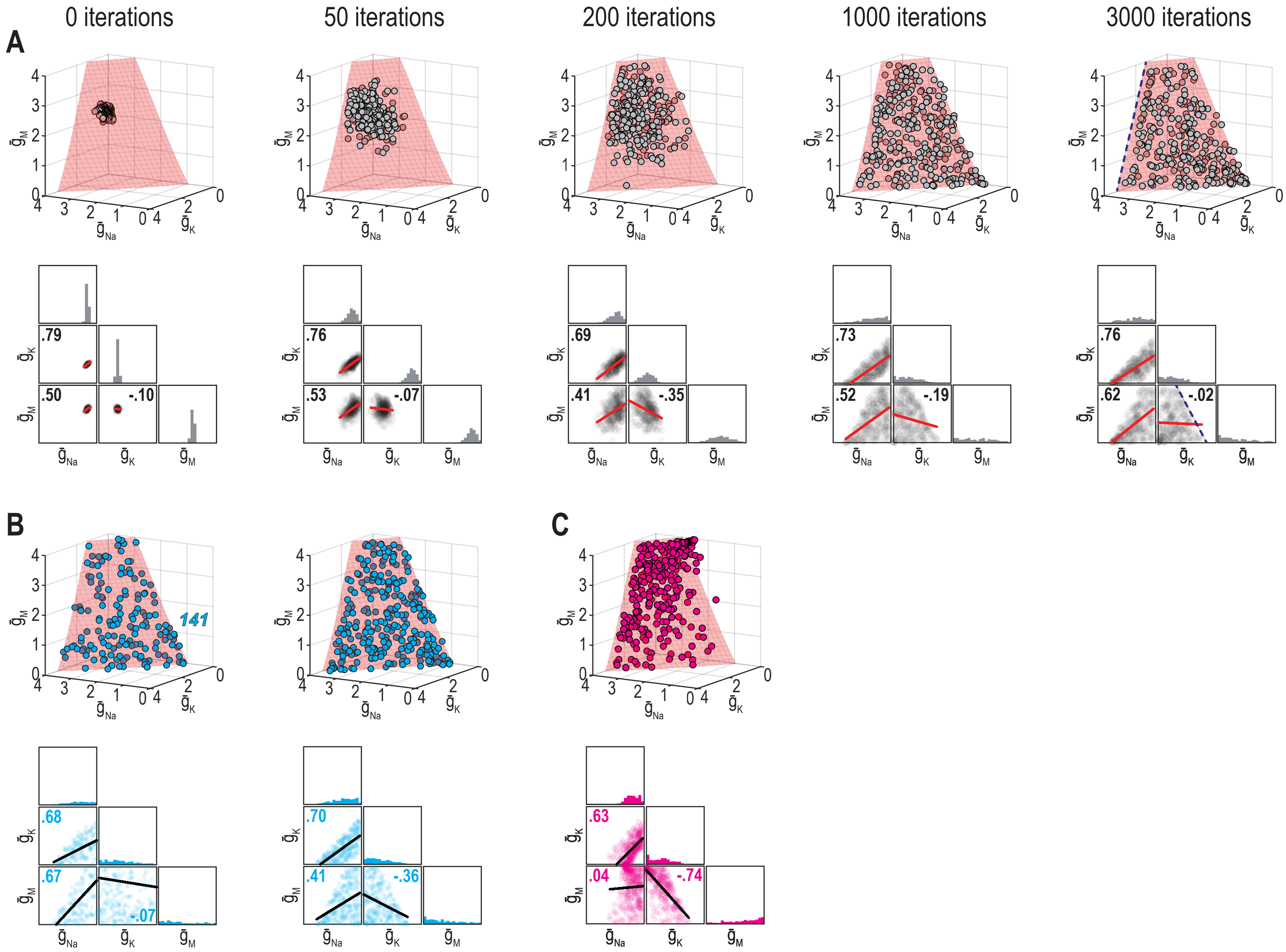


**Figure 8 - figure supplement 1**

**Table S1. Initial conductance values and regulation rates for each figure.**

**Initial conductance values ($mS/cm^2$):**

| | $\bar{g}_{Na}$ | $\bar{g}_K$ | $\bar{g}_{leak}$ | $\bar{g}_M$ | $\bar{g}_{AHP}$ |
|---|---|---|---|---|---|
| **mean** | **3.5** | **0.7** | **2** | **1.75** | **0.5** |
| **stdev** | **0.1** | **0.1** | **0.1** | **0.1** | **0.1** |

**Regulation rates $\tau_{i,j}$:**

| | **ion channel i** | | | | | |
|---|---|---|---|---|---|---|
| | $\bar{g}_{Na}$ | $\bar{g}_K$ | $\bar{g}_{leak}$ | $\bar{g}_M$ | $\bar{g}_{AHP}$ | **error signal j** |
| **Figure 4** | -60 | N/A | 60 | N/A | N/A | FR |
| **Figure6** | -400 | 100 | N/A | 100 | 100 | FR |
| | 0.15 | 0.105 | N/A | 0.105 | 0.105 | EE |
| **Figure 7 - cyan** | -300 | 300 | N/A | 60 | N/A | FR |
| | 0.25 | -0.05 | N/A | 0.025 | N/A | EE |
| **Figure 7 - magenta** | -360 | 60 | N/A | -120 | N/A | FR |
| | 0.2143 | -0.05 | N/A | 0.0714 | N/A | EE |
| **Figure 9** | -100 | -600 | N/A | -180 | N/A | FR |
| | 0.15 | -0.09 | N/A | -0.06 | N/A | EE |
| **Figure 10** | -105 | 225 | N/A | -210 | N/A | FR |
| | 0.0625 | -0.625 | N/A | 0.0875 | N/A | EE |

Firing rate (FR); energy efficiency (EE).
